# Multilayer interaction networks reveal multidimensional ecological strategies and context-dependent competition in soil actinobacteria

**DOI:** 10.64898/2026.09.16.752070

**Authors:** Youssef Zerta, Jaouad Anissi

**Affiliations:** Euromed University of Fes, Route Nationale Fès-Meknès, Fez, Morocco

**Keywords:** actinobacteria, *Streptomyces*, multilayer interaction network, pairwise interaction phenotypes, ecological strategy, spatial dependence

## Abstract

Microbial communities are structured by interactions spanning antagonism, inducible responses, metabolic modulation, and morphogenetic effects, yet these processes are often reduced to a single antagonistic dimension. Here, we characterized pairwise interactions among 60 soil-derived actinobacterial strains, predominantly Streptomyces, across 12 phenotypes represented as a binary, directed multilayer network. The layers were neither reducible to a single ecological axis nor globally independent. Degree-preserving null models identified significant excess overlap in 8 of 65 layer pairs. Although antimicrobial release and growth inhibition formed the principal connectivity backbone, strongly connected component sizes were consistent with degree-based null expectations. Strain-level strategies formed a heterogeneous multidimensional space. Two principal components exceeded the 95th percentile of Horn’s parallel-analysis null, explaining 49.8% of variance; a third axis exceeded only the weaker mean-eigenvalue criterion and is reported as marginal. Gaussian mixture modelling gave no stable component count across covariance parameterizations, leaving the number of ecological groups unresolved rather than excluded. Dominance was phenotype dependent, with weak cross-layer correlations of David’s scores and no universal sender. Interaction responses also showed phenotype-specific spatial dependence. Ecological strategy was unrelated to 16S rRNA phylogeny. Pagel’s λ showed no significant signal across 21 traits (all *q* ≥ 0.39), and multilayer ecological distance was unrelated to 16S rRNA distance (*ρ=*-0.04, *P=*0.59), although inference was constrained by marker resolution. These results reveal a heterogeneous, multidimensional interaction landscape structured by partially coupled functional phenotypes rather than a single antagonistic hierarchy, potentially reflecting semi-independent evolution of secondary metabolism, development, sensing and stress-response pathways.

## 1. Introduction

Microbial communities are one of the most complex and dynamic biological systems on Earth, shaped by intricate networks of interactions that span competition, cooperation, signalling, and environmental adaptation [1]. Since Darwin’s early reflections on the struggle for existence [2], the idea that coexistence emerges from a balance of opposing forces has remained central to ecological theory [3]. In microbial systems, this balance is particularly striking: despite intense competition for limited resources, highly diverse communities persist in soils, aquatic environments, and host-associated microbiomes [4, 5]. Understanding how such diversity is maintained requires not only identifying the interacting species but also deciphering the mechanisms and structures of their interactions.

Among soil microorganisms, *Streptomyces* species occupy a unique ecological and evolutionary position [6]. These filamentous, Gram-positive bacteria are prolific producers of secondary metabolites, including antibiotics, antifungals, and signalling molecules, and play a critical role in nutrient cycling and organic matter decomposition [7]. Their genomes harbour large numbers of biosynthetic gene clusters (BGCs), enabling them to produce a wide array of diffusible and volatile compounds that mediate interactions with neighbouring organisms [8-10]. As a result, *Streptomyces spp.* are often viewed as central players in microbial chemical warfare, shaping community composition through inhibition, resistance and modulation mechanisms [11]. However, this view, where interactions are primarily framed as antagonistic, has increasingly been challenged [12]. Recent studies suggest that microbial communities are structured not only by competition but also by facilitation, metabolic interdependence, and regulatory signalling [13]. For instance, antibiotic production can induce responses in neighbouring cells, triggering defence mechanisms or even cooperative behaviours [14, 15]. Similarly, metabolic by-products can serve as nutrients for other species, creating networks of cross-feeding [16]. Here, we investigated whether the diversity of pairwise interaction phenotypes among soil-derived actinobacteria reflects a common antagonistic organization or a multidimensional ecological structure. We asked whether the twelve phenotypes differ in network architecture and overlap more than their degree sequences predict, whether strain-level strategies follow one influence ranking or vary across several dimensions, whether two of these phenotypes depend on the spatial configuration of the confrontation, and whether interaction profiles track relatedness at the resolution near-full-length 16S rRNA provides.

## 2. Materials and Methods

### 2.1. Bacterial isolation and strain characterization

Sixty soil-derived actinobacterial strains were investigated, predominantly belonging to the genus *Streptomyces*. They originated from soils collected across the Rif and Atlas mountain regions of Morocco. Soil suspensions were serially diluted and cultured on Bennett’s agar, and purified isolates were maintained under standardized conditions. Taxonomic characterization was based on near-full-length 16S rRNA gene sequences obtained by Sanger sequencing. Sequences were aligned with MAFFT (FFT-NS-i) [17], trimmed with trimAl (gappyout; 1,533 to 1,337 columns) [18], and a maximum-likelihood tree was inferred under GTR+GAMMA with VeryFastTree, a FastTree 2 implementation [19, 20]. Biochemical characterization was performed using API 50CH, API 20 NE and API ZYM systems (BioMérieux, France). Strain identities and sequence accession numbers are provided in Supplementary Table S1; sensitivity to excluding the four non-*Streptomyces* isolates is reported in Supplementary Table S4. Detailed procedures are provided in Supplementary Methods.

### 2.2. Pairwise interaction assay and phenotype scoring

Pairwise interactions were evaluated on Bennett’s agar for all ordered strain pairs (*S_i_,S_j_*). Under direct contact, 10 µL spore suspensions (∼2.4*10^6^ CFU/mL) were inoculated close enough to permit physical contact and scored after 5-6 days at 30 °C for twelve phenotypic responses: CMP, CS, CCVM, CCAM, RP, IRP, IG, IC, RAC, IAC_RAC, IRAC and IAC_RDE. The four antimicrobial phenotypes used a soft-agar overlay with Bacillus subtilis ATCC 6051 as indicator, following Davelos *et al.* (2004) [21]. The complete set of confrontations was performed in two independent runs, and each plate was read twice by the same investigator with review by a second investigator; runs and readings agreed except for rare discordant calls, which were resolved by re-reading the plate before a single binary score was recorded. IG and CS were additionally evaluated under spatial separation, analysed separately rather than as a further network dimension. Spatial configurations and paired statistical analyses are described in Supplementary Methods S10. Full definitions and scoring criteria are in Supplementary Methods S2. Several layers share an assay context, which is treated as a limit on interpretation throughout (Supplementary Methods S4).

### 2.3. Multilayer network representation

Each direct-contact phenotype was represented as a directed binary 60x60 adjacency matrix, in which an edge from strain *S_i_*} to strain *S_j_* indicated that *S_i_* elicited the corresponding phenotype in *S_j_*. Thus, the twelve matrices collectively defined a directed multilayer interaction network. Two layers qualify this definition. RP was scored as a change in the pigmentation of *S*_j_ relative to its own monoculture control, and is directed in that sense. RAC was read from a soft-agar overlay against the indicator strain and is dominated by the strain in which it is scored rather than by the pairing, so its entries record largely whether that strain releases antibacterial compounds at all. It is retained as a layer throughout, with that constraint carried wherever its layer-level statistics are reported.

For biological interpretation, layers were additionally grouped into four functional categories: direct antagonism DA={IG, IC, RAC}, induced antagonism IA={IAC_RAC, IAC_RDE, IRAC}, metabolic/pigment modulation MM={RP, IRP}, and morphogenetic/developmental control MC={CCAM, CCVM, CMP, CS}. These categories were used to construct a complementary four-dimensional ecological representation of strain interaction profiles.

### 2.4. Network structure and cross-layer organization

For each interaction layer, we quantified directed edge count and density, pair-based reciprocity, transitivity on the undirected projection, out-degree, and the size of the largest strongly connected component (SCC). Self-interactions were excluded throughout, leaving 3,540 possible directed edges for *N*=60 strains. Reciprocity is the proportion of connected dyads reciprocated in both directions, and a participation coefficient [22] quantified whether a strain’s outgoing interactions concentrated in particular phenotypes or spread across layers. Cross-layer similarity used the Jaccard index [23] on directed edge sets. Each layer was randomized into 10,000 degree-preserving null networks by directed edge swaps [24] that retain its empirical in- and out-degree sequences, and the same draws served as the null for overlap, SCC size, reciprocity and transitivity alike. Two-sided empirical P-values used the Phipson-Smyth correction [25], with Benjamini-Yekutieli FDR control [26] applied separately within each family of tests. The CCAM-CCVM pair was designated a priori as a shared-measurement confound, both colours being scored from the same colonies, and is reported separately from the biological summary (Supplementary Methods S4).(*N* (*N −* 1)=3540) (*R_p_*=*M* / (*M* +*U*))

### 2.5. Multidimensional ecological strategies

Strain profiles were the z-standardized twelve-layer out-degree vectors. Non-trivial PCA axes were assessed using permutation-based Horn’s parallel analysis [27, 28], comparing observed variances with 1,000 independently permuted datasets. Components were retained when the observed variance exceeded the 95th percentile of the corresponding null distribution; the weaker mean-eigenvalue criterion is reported alongside as a sensitivity analysis. Discrete grouping was evaluated using Gaussian mixture models with *k=*1-8 components on 12-dimensional profiles and four-dimensional DA-IA-MM-MC representations. Models were compared using BIC, ICL, and 10-fold cross-validated held-out log-likelihood [29, 30]. Given *N=*60 and full-covariance model complexity, clustering required consistent model-selection evidence, with disagreements interpreted conservatively [31].

Univariate multimodality along each retained principal component was assessed using Hartigan’s dip test [32]. Because unimodality of individual components does not imply unimodality of the joint distribution, dip results were treated as complementary evidence only.

### 2.6. Layer-specific ecological influence

Ecological influence was evaluated independently within each interaction layer using normalized David’s scores [33, 34]. Bradley-Terry models [35] were fitted to the same matrices as a complementary summary of directional asymmetry; because both estimators derive from the same data, their agreement measures methodological concordance rather than independent validation. To determine whether ecological influence represented a general strain property or was phenotype specific, pairwise Spearman correlations were calculated among layer-specific David’s scores, followed by PCA of the 60*12 dominance matrix. Two multilayer descriptors summarize how stable a strain’s direction of influence is across phenotypes. Within each layer, a strain was classed as a net sender when its out-degree exceeded its in-degree and as a net receiver in the converse case, layers in which the two were equal being left unclassified. Role instability, referred to as sender/receiver switching, is the number of classified layers whose class differs from that strain’s majority class, from 0 for one role throughout to 6 for an even split. Variability of interaction output is summarized separately as the standard deviation of a strain’s out-degree across the twelve layers. A consensus interaction network was additionally constructed by retaining directed interactions occurring in at least four of the twelve phenotypic layers to identify recurrent multilayer relationships.

### 2.7. Association between 16S rRNA relatedness and interaction profiles

Phylogenetic analyses were conducted for all 60 isolates. One isolate appears as S1430 in the sequencing record and as S1226 in the interaction matrices; the two labels denote the same strain. Phylogenetic signal was evaluated for 21 strain-level ecological traits, comprising the twelve layer-specific out-degrees, four functional-category scores, total out-degree, participation coefficient and the first three PCA axes; PC3 is included as a descriptive axis although it did not meet the 95th-percentile retention criterion. Pagel’s λ was estimated by maximum likelihood [36]. Because identical 16S rRNA sequences produced a rank-deficient phylogenetic covariance matrix and conventional likelihood-ratio inference was anti-conservative (Fig. S9), significance was determined using 999 tip-label permutations followed by Benjamini-Hochberg correction [37]. Robustness was evaluated using alternative tree reconstructions and a tree-independent Mantel test [38] relating pairwise 16S rRNA distance to ecological distance, defined as the Euclidean distance between z-standardized twelve-layer out-degree profiles; because the 1,770 dyadic distances are not independent, Mantel significance was assessed by 9,999 permutations of strain labels. These analyses test for structure at the resolution of 16S rRNA only and do not address fine-scale evolutionary independence.

### 2.8. Data availability

Binary interaction matrices, including the paired spatial matrices, 16S rRNA records and all analysis code are available at https://doi.org/10.6084/m9.figshare.33826675. 16S rRNA sequences are deposited in GenBank, with accession numbers provided in Supplementary Table S1.

## 3. Results

### 3.1. Direct*-*contact assays reveal a multidimensional repertoire of ecological interaction phenotypes

Pairwise interaction assays under direct-contact conditions revealed extensive phenotypic diversity across the collection, demonstrating that strain-strain interactions were not restricted to growth inhibition but encompassed several functionally distinct responses. Individual strain combinations generated markedly different outcomes under the same culture conditions (Fig. 1; Fig. S1), which is the basis for representing the system as a directed multilayer network.

**Figure 1:**
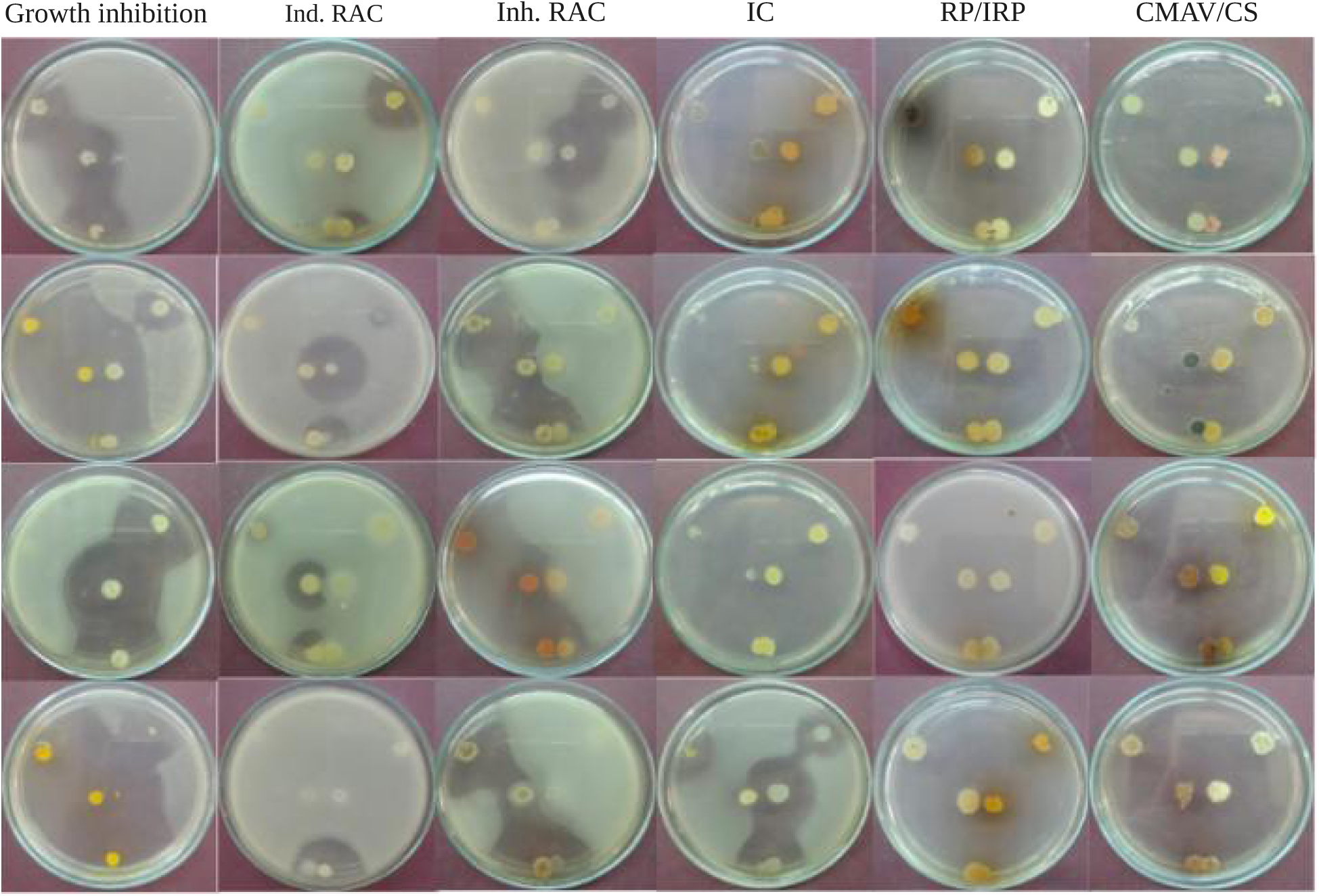
Representative interaction phenotypes among Streptomyces strains. Representative examples of direct and indirect ecological interactions observed during pairwise confrontation assays. Shown are inhibition of growth (IG), induced antagonistic responses (IRAC), release of degrading enzymes (RDE), invasion competition (IC), induced pigment production (IPP), and colony coloration or morphogenetic changes (CCVM). Together, these examples illustrate the diversity of ecological mechanisms operating within Streptomyces communities and forming the basis of the multilayer interaction framework.

The twelve direct-contact layers differed strongly in their prevalence and distribution among strain pairs, revealing substantial heterogeneity in both sender and receiver behaviour (Fig. S1). Direct-antagonism phenotypes were particularly widespread. For example, antimicrobial release (RAC) formed the densest layer, with 1,677 directed edges (density 0.474) and pair-based reciprocity of 0.297, followed by growth inhibition (IG; 767 edges, density 0.217, reciprocity 0.150). Colony invasion (IC), although also classified as direct antagonism, was considerably sparser and predominantly asymmetric. Among inducible-antagonism phenotypes, IAC_RAC and IRAC exhibited intermediate connectivity, whereas IAC_RDE occurred comparatively rarely. Developmental and morphogenetic responses, including CS, CMP, CCVM, and CCAM, displayed intermediate but strongly strain-dependent patterns, while RP and IRP represented among the sparsest interaction dimensions (Fig. S1). No single phenotype therefore captured the full repertoire of pairwise responses. We next asked whether these differences reflected distinct network architectures or only differences in density.

### 3.2. Interaction layers are weakly redundant but exhibit functionally coherent overlap

Raw edge overlap was low across interaction layers (mean Jaccard similarity = 0.047; 0.043 excluding the prespecified CCAM-CCVM confound; Fig. S2). Degree-preserving null models showed that most layer pairs matched degree-constrained expectations, whereas nine of 65 interpretable comparisons deviated significantly: eight were enriched and one depleted (Fig. 2; Table 2; Supplementary Table S2). Significant deviations were concentrated among developmental/morphological and antimicrobial-related phenotypes, indicating selective cross-layer correspondence. Because several phenotypes shared assay contexts, enriched overlap cannot be attributed solely to biological coupling and was interpreted as non-random phenotypic correspondence rather than evidence of shared regulatory mechanisms.

**Figure 2.**
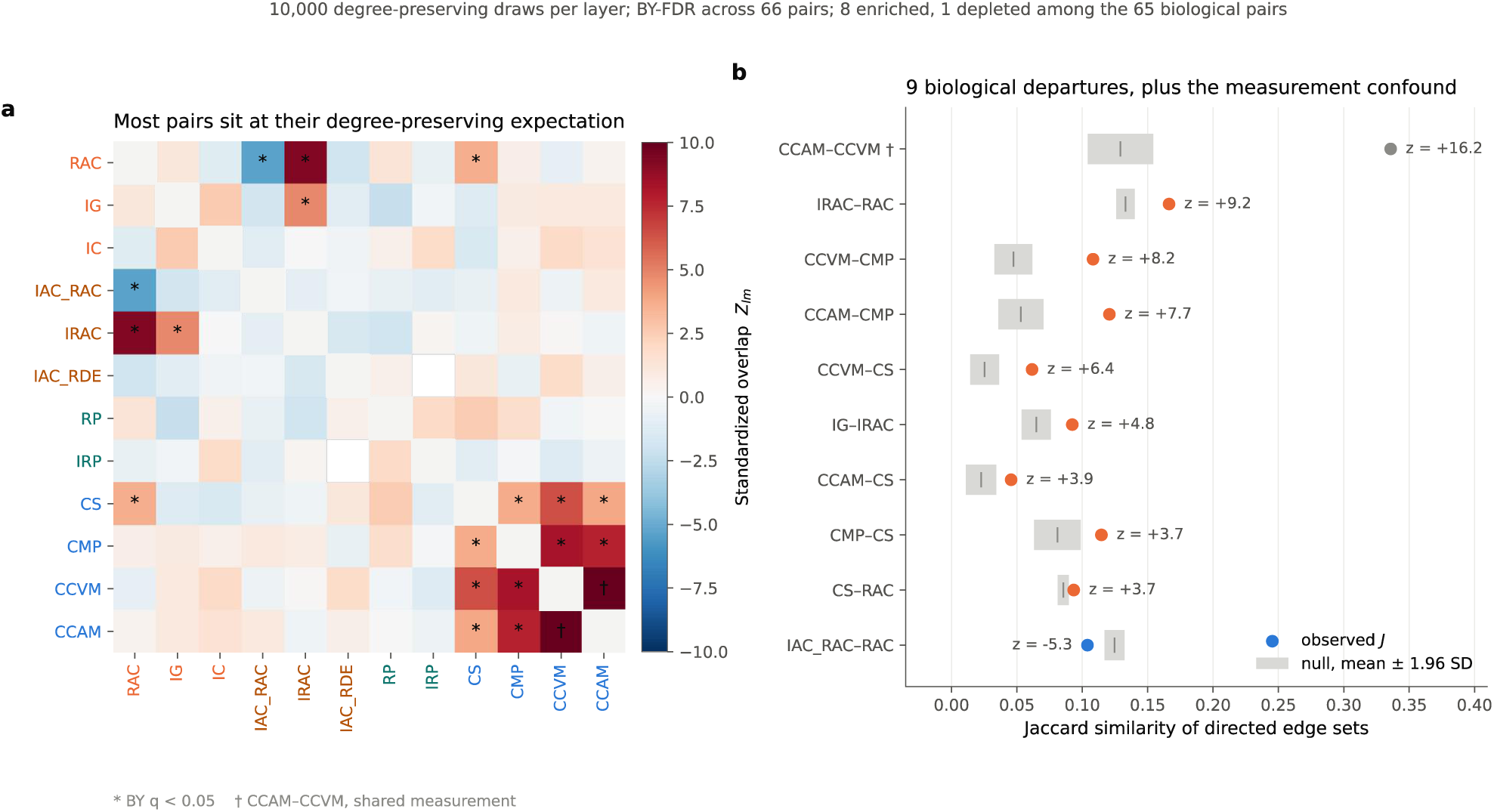
Selective cross-layer correspondence relative to degree-preserving null expectations. a, standardized overlap Zlm for all 66 layer pairs. For each pair the observed Jaccard similarity of directed edge sets was compared with a null from 10,000 degree-preserving directed edge-swap randomizations of each layer, *Z* =(*Jobs – mean* (*Jnull*))/ *sd* (*Jnull*); positive values indicate more shared edges than the degree sequences predict. Asterisks mark Benjamini-Yekutieli *q* < 0.05 across all 66 pairs. b, the ten pairs that depart from the null, showing observed Jaccard similarity against the null mean ± 1.96 SD. Nine are biological departures, eight enriched and one depleted; CCAM-CCVM is marked † because both colours are scored from the same colonies, so shared measurement and shared biology cannot be separated for that pair. The degree-preserving null controls for density and degree structure but not for covariance introduced by shared assays, so enriched overlap is evidence of non-random phenotypic correspondence rather than of common regulation.

### 3.3. Direct antagonism dominates network connectivity, whereas null models reveal phenotype*-*specific higher*-*order organization

Network architecture varied markedly among phenotypes (Table 1; Fig. S3). RAC was the densest layer (0.474), followed by IG (0.217), IAC_RAC (0.108), and IRAC (0.091), whereas developmental, morphogenetic, and metabolic layers were substantially sparser. Reciprocity was highest in RAC (0.297) and IG (0.150), remained below 0.10 in all other layers, and was absent in IRP and IAC_RDE. IG contained the largest strongly connected component (SCC; 59 strains), followed by RAC (51). However, degree-preserving null models showed that SCC sizes remained within null expectations across all layers (Fig. S3), indicating that broad connectedness largely reflected empirical degree distributions. Five layers were significantly more reciprocal than expected after Benjamini-Yekutieli correction: IG (*z=*+6.17), IRAC (+4.24), CS (+3.10), IAC_RAC (+2.82), and RAC (+2.75). No layer was significantly less reciprocal, and reciprocity was undefined for IRP and IAC_RDE. Conversely, four layers were significantly less transitive than expected: CS (*z=*-4.06), RAC (-4.05), IG (-3.56), and CMP (-3.34), while CCVM approached significance (*z*=-2.60, *q=*0.064) (Fig. S3). RAC, despite its high density and reciprocity, was among the least transitive. Because RAC was dominated by the scored strain (57.6% of deviance versus 0.8% for its partner), its statistics mainly reflect broad antibacterial release rather than partner-specific organization. In contrast, IG retained a genuinely dyadic structure. Thus, direct antagonism forms the principal connectivity backbone, while reciprocity, SCC structure, and transitivity reveal distinct phenotype-specific organization (Fig. S3).

**Table 1.**
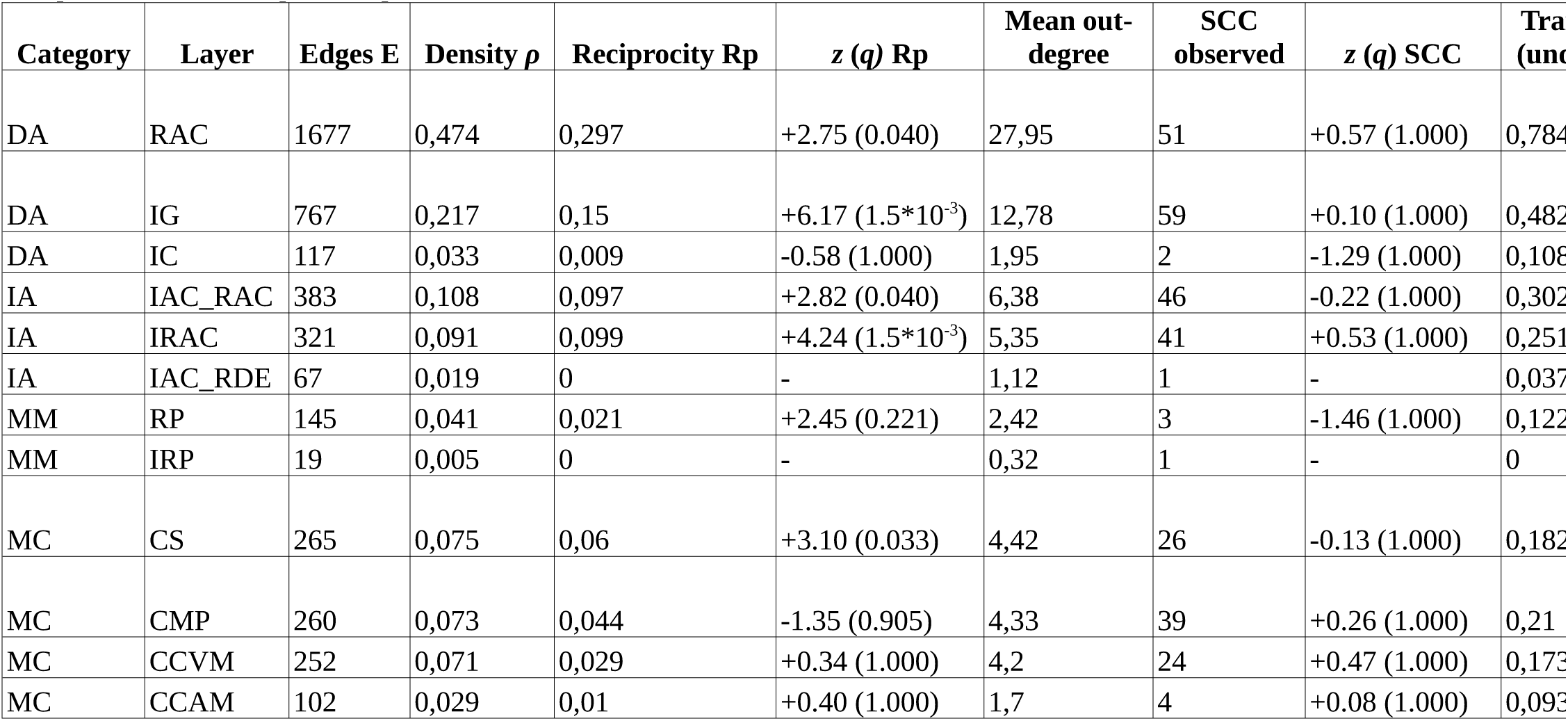
Per-layer architecture of the twelve interaction networks. *N*=60 strains, binary directed projection, self-interactions excluded (*N* (*N −* 1)=3,540 possible directed edges). Density ρ=E / N(N-1) with N(N-1)=3,540 ordered non-self pairs; pair-based reciprocity Rp=mutual / connected dyads; SCC=size of the largest strongly connected component; transitivity is computed on the undirected projection of each layer. Each *z* compares the observed value with 10,000 degree-preserving directed edge-swap rewirings retaining the empirical in- and out-degree sequences, *z*=(observed null mean) / null SD; the value in brackets is the Benjamini-Yekutieli-corrected empirical probability within that statistic’s family of twelve layers. An em-dash marks a degenerate null in which the statistic is identical in every draw. RAC is dominated by the strain in which it is scored, whose identity accounts for 57.6% of the deviance in that matrix against 0.8% for its partner, so its layer-level values summarise how broadly strains release antibacterial compounds rather than partner-specific structure.

**Table 2.**
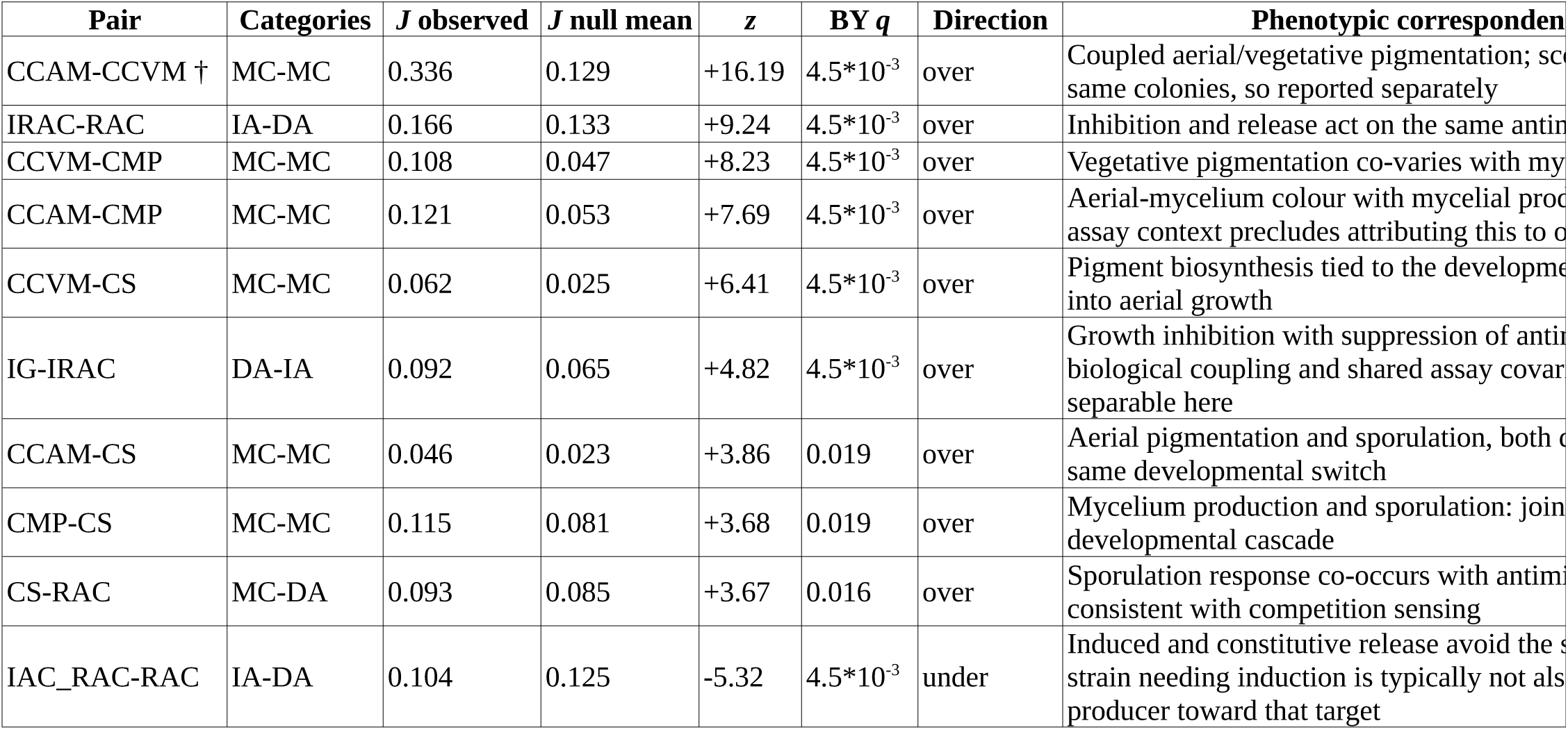
Layer pairs deviating from a degree-preserving null. Jaccard similarity Jobs on directed edge sets compared with 10,000 independent degree-preserving edge-swap rewirings of each layer; *z*=(*Jobs −mean* (*Jnull*))/ *sd* (*Jnull*). Two-sided empirical probabilities use the Phipson-Smyth (*M* +1) / (*N* +1) correction and are Benjamini-Yekutieli-corrected across all 66 pairs, BY rather than BH because each layer appears in eleven tests. Only pairs with *q* < 0.05 are listed; Table S2 gives all 66. The final column names the phenotypic correspondence observed. It is deliberately not a mechanistic account: these are binary co-occurrence data with no molecular measurements, and several of these pairs share an assay context, so a shared regulatory explanation is not identifiable from this design. † CCAM-CCVM is a prespecified shared-measurement pair, reported separately from the biological summary.

| Pair | Categories | J observed | J null mean | z | BY q | Direction | Phenotypic correspondence |
| --- | --- | --- | --- | --- | --- | --- | --- |
| CCAM-CCVM † | MC-MC | 0.336 | 0.129 | +16.19 | $4.5 \times 10^{-3}$ | over | Coupled aerial/vegetative pigmentation; same colonies, so reported separately |
| IRAC-RAC | IA-DA | 0.166 | 0.133 | +9.24 | $4.5 \times 10^{-3}$ | over | Inhibition and release act on the same antimicrobial target |
| CCVM-CMP | MC-MC | 0.108 | 0.047 | +8.23 | $4.5 \times 10^{-3}$ | over | Vegetative pigmentation co-varies with mycelium production |
| CCAM-CMP | MC-MC | 0.121 | 0.053 | +7.69 | $4.5 \times 10^{-3}$ | over | Aerial-mycelium colour with mycelial production; assay context precludes attributing this to one |
| CCVM-CS | MC-MC | 0.062 | 0.025 | +6.41 | $4.5 \times 10^{-3}$ | over | Pigment biosynthesis tied to the developmental switch into aerial growth |
| IG-IRAC | DA-IA | 0.092 | 0.065 | +4.82 | $4.5 \times 10^{-3}$ | over | Growth inhibition with suppression of antimicrobial release; biological coupling and shared assay covary, but not separable here |
| CCAM-CS | MC-MC | 0.046 | 0.023 | +3.86 | 0.019 | over | Aerial pigmentation and sporulation, both controlled by the same developmental switch |
| CMP-CS | MC-MC | 0.115 | 0.081 | +3.68 | 0.019 | over | Mycelium production and sporulation: joint control by a developmental cascade |
| CS-RAC | MC-DA | 0.093 | 0.085 | +3.67 | 0.016 | over | Sporulation response co-occurs with antimicrobial release; consistent with competition sensing |
| IAC_RAC-RAC | IA-DA | 0.104 | 0.125 | -5.32 | $4.5 \times 10^{-3}$ | under | Induced and constitutive release avoid the same target; strain needing induction is typically not also a producer toward that target |

### 3.4. Strain*-*level interaction strategies occupy a multidimensional rather than a single antagonistic axis

The weak redundancy among interaction layers suggested that strain-level interaction profiles required multiple dimensions for adequate representation. Principal component analysis of the 12-dimensional outgoing profiles showed that PC1 and PC2 explained 32.5% and 17.3% of total variance, respectively (49.8% cumulatively), and both exceeded the 95th percentile of the Horn parallel-analysis null (17.0% and 14.4% respectively; Fig. S4). PC3 explained a further 11.8%, exceeding the mean null eigenvalue (11.6%) but not its 95th percentile (12.6%), with a bootstrap interval (9.8-15.0%) that spans the null; PC3 is therefore reported as marginal rather than as an established axis. PC4 (8.2%) fell below the null on both criteria. PC1 integrated developmental, pigmentation-associated, and inhibitory responses, with strongest contributions from CMP (0.41), IRP (0.40), CCVM (0.38), CCAM (0.37), and IG (0.33). PC2 differentiated direct-antagonism phenotypes, with RAC loading positively (0.54), whereas IC and IG contributed oppositely (approximately -0.38), indicating that direct-antagonism traits did not vary coordinately. PC3 was dominated by IAC_RDE (0.65), with additional contributions from IAC_RAC (-0.39) and RP (0.37). Gaussian mixture models fitted to the 12-dimensional profiles and the four-dimensional functional-category representation gave conflicting evidence. BIC and ICL favoured multi-component solutions, the preferred k ranging from four to eight across embeddings and covariance structures, whereas cross-validated held-out likelihood favoured one component under full covariance and two to three under the constrained parametrizations, the diagonal three-component model scoring highest of any model examined. Given *N=*60 and the free parameters of the unconstrained models, this disagreement leaves the number of components unresolved rather than absent. Together with the broad spread of strains across the retained axes, interaction profiles are better described as heterogeneous and multidimensional than as separated ecological groups (Fig. S5; Fig. 3).

**Figure 3.**
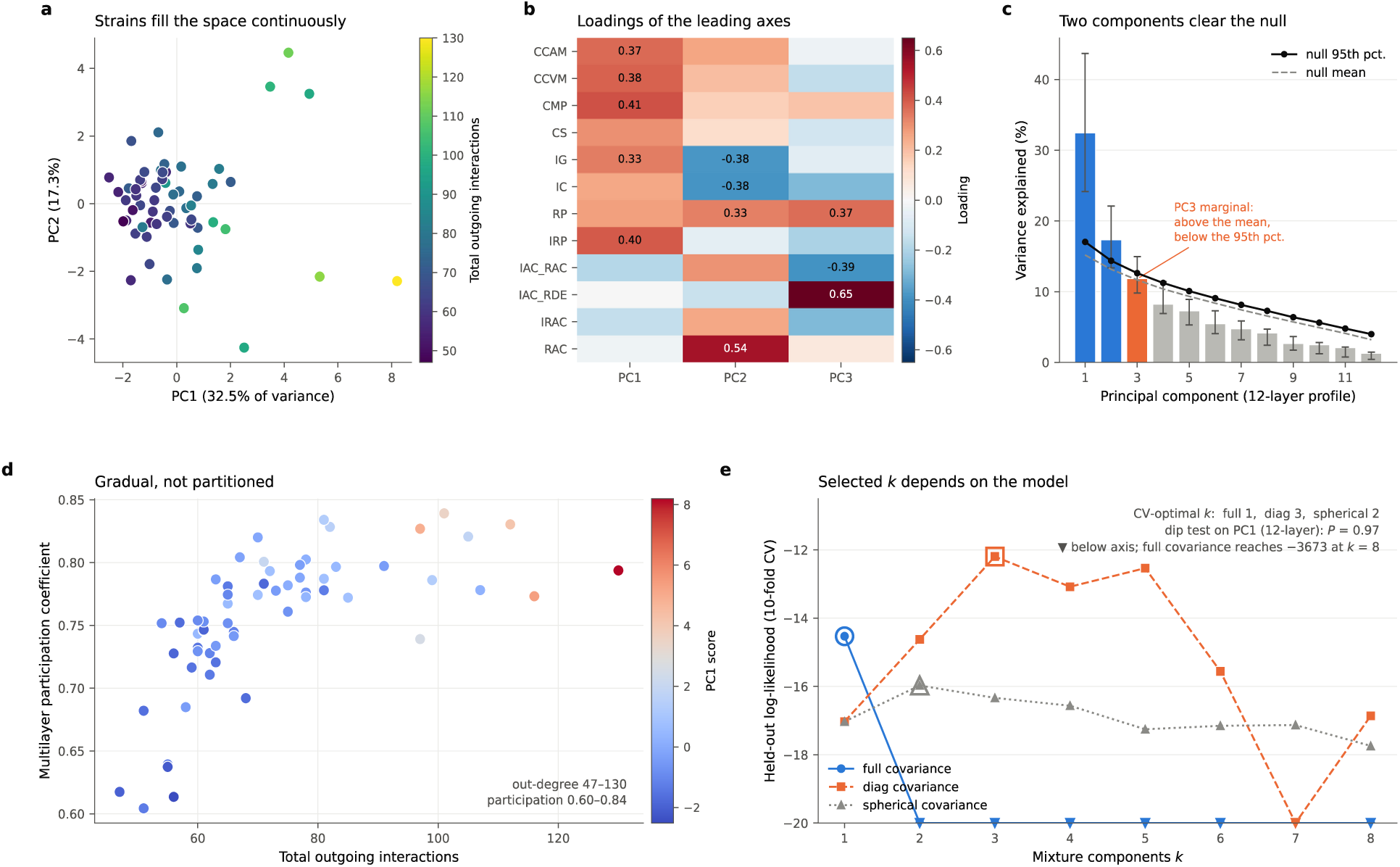
Strain interaction strategies occupy a multidimensional space of unresolved cluster structure. a, principal component analysis of the twelve-layer outgoing profile on z-standardized counts, coloured by total outgoing interactions. b, loadings of the three leading axes on each layer; values of magnitude ≥ 0.30 are printed. c, variance explained by each component against the Horn parallel-analysis null from 1,000 column-permuted datasets, with 95% bootstrap intervals from 2,000 resamples of the strains. PC1 (32.5%) and PC2 (17.3%) exceed the 95th percentile of the null; PC3 (11.8%) exceeds the null mean (11.6%) but not its 95th percentile (12.6%) and is drawn in orange as marginal. d, total outgoing interactions against multilayer participation for all 60 strains; out-degree spans 47-130 and participation 0.60-0.84, and the distribution is gradual rather than partitioned into generalists and specialists. e, held-out log-likelihood by component count under three covariance structures. The selected *k* depends on the model, full covariance selects *k*=1, diagonal *k*=3 and spherical *k*=2, and the diagonal three-component solution attains the highest held-out likelihood of any model examined, so the number of components is unresolved rather than established at one.

### 3.5. Multilayer participation reveals a continuum from specialized to broadly interactive strains

If ecological strategies are multidimensional, strain-level influence should vary across phenotypes rather than remain fixed among interaction layers. Total outgoing degree varied markedly, from 47 to 130 interactions, while most strains showed relatively high participation coefficients (0.60-0.84), so activity is distributed across multiple layers. The most active strains combined high output with broad participation across all four functional categories, while others paired substantial output with lower participation, indicating specialization within fewer layers (Fig. S6). The distribution was gradual rather than partitioned into discrete generalists and specialists (Fig. 3d). Aggregating the twelve layers into the four functional dimensions preserved much of this ordering, but because DA, IA, MM and MC were defined a priori from those same layers, this is an interpretable compression rather than independent validation. The first principal axis of the reduced four-dimensional representation explained 53.0% of the variance and was strongly correlated with PC1 of the full twelve-layer profile (Spearman *ρ=*0.934, *P=*1.5*10^-27^; Fig. S7). Hartigan’s dip test on the first axis of this four-dimensional representation was consistent with unimodality (*P=*0.531; the corresponding test on PC1 of the full twelve-layer profile gave *P=*0.967), supporting that interaction profiles varied across a multidimensional strategy space, with no robust statistical evidence for discrete ecological groups at the present sample size (Fig. 3).

### 3.6. Ecological influence is phenotype dependent and recurrent interactions are concentrated among highly interactive strains

If strains occupied fixed dominance positions across the multilayer network, their sender-receiver rankings should remain consistent among phenotypes. Instead, extensive rank rearrangements occurred across layers, with individual strains shifting between strong sender and receiver positions depending on the phenotype considered. Accordingly, correspondence among layer-specific David’s scores was weak. Across off-diagonal comparisons, mean and median Spearman correlations were only *ρ=*0.063 and *ρ=*0.043, respectively. The strongest positive correlation involved the co-scored CCAM-CCVM pair (*ρ=*0.62); among the other layer pairs the strongest were IRAC-RAC (*ρ=*0.59) and IG-IC (*ρ=*0.56). The strongest association of any sign, however, was negative: David’s scores in IG and in RAC were inversely related (*ρ=*-0.66). This does not reflect distinct capacities. RAC is scored on the releasing strain, which occupies the receiver position of the matrix convention, so a strain’s RAC David’s score falls as the number of partners against which it releases antibacterial compounds rises (*ρ=*-0.97); read through that count, the strains that most often inhibited a partner’s growth were also those that most often released antibacterial compounds (Spearman *ρ=*0.75 between IG out-degree and RAC release count), and the RAC score is therefore not read as sender influence. Consistently, six principal components were required to explain ≥80% of dominance variance, with cumulative variance reaching 24.3%, 41.2%, 55.3%, 67.6%, 77.6%, and 84.5% across PC1-PC6. The agreement between the two estimators supports methodological concordance within layers: Bradley-Terry strengths and David’s scores were strongly concordant (Spearman *ρ=*0.59-0.99; mean 0.87; Table S3). No single strain-level ranking therefore summarizes the complete interaction repertoire. For explicitly antagonistic layers, differences in sender influence can additionally be interpreted in terms of competitive effects; however, such a dominance interpretation was not extended to developmental, morphological, metabolic, or induced-response phenotypes (Fig. 4; Table S3).

**Figure 4.**
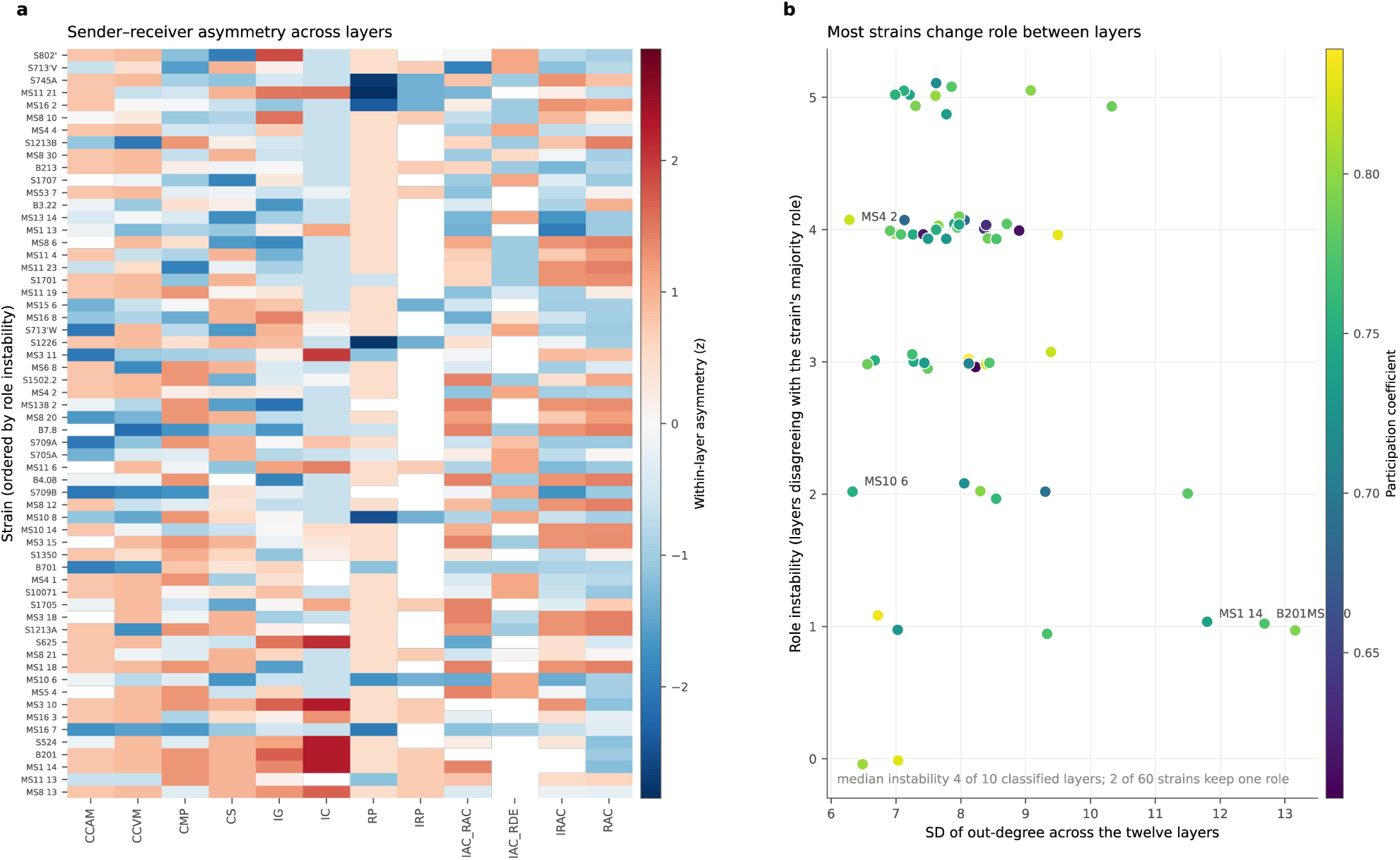
Directional influence is phenotype dependent rather than a fixed strain property. a, within-strain sender-versus-receiver asymmetry, *z*-scored within each layer, for all 60 strains across the twelve layers; strains are ordered by role instability. White cells mark layers in which a strain neither sends nor receives. b, role instability against the variability of interaction output, coloured by participation coefficient. Within a layer a strain is a net sender when its out-degree exceeds its in-degree and a net receiver when the reverse holds, with ties left unclassified; role instability is the number of classified layers whose class differs from that strain’s own majority class, so 0 marks a strain holding one role throughout and 6 an even split. The measure does not depend on the ordering of the layers. Correspondence among layer-specific David’s scores is weak (mean Spearman *ρ*=0.063, median 0.043) and six principal components are needed to explain ≥80% of dominance variance, so no single ranking summarises the repertoire. Within layers, Bradley-Terry strengths and David’s scores agree closely (*ρ*=0.59-0.99, mean 0.87; Table S3), so the two estimators are concordant, though both derive from the same matrices and so do not validate one another independently.

We next examined whether particular interactions nevertheless recurred across phenotypes by constructing a consensus network retaining directed edges present in at least four of the twelve layers (Fig. S8). Recurrent interactions were distributed heterogeneously, with a relatively small subset of strains participating in numerous consensus edges while many others remained weakly connected or peripheral. Strains previously characterized by high total out-degree and broad participation were frequently represented among these recurrently connected nodes (Fig. S8a). Because the ≥4-layer criterion is operational rather than biologically derived, this indicates heterogeneous multilayer recurrence rather than core-periphery organization.

### 3.7. No detectable association of interaction profiles with 16S relatedness or measured metabolic niche

We next tested whether strain-level interaction variation was associated with relatedness as resolved by near-full-length 16S rRNA sequences. Across the 21 ecological traits examined, observed Pagel’s λ values remained within their respective tip-label permutation distributions after multiple-testing correction (minimum *q=*0.39; Fig. 5a). Consistently, pairwise 16S rRNA genetic distance was unrelated to ecological distance calculated from the complete 12-layer interaction profiles (Mantel *ρ=*-0.041, *P=*0.59, 9,999 strain-label permutations; Fig. 5b). Layer-specific analyses using Hamming and Jaccard ecological distances likewise provided no significant evidence of association with 16S rRNA divergence (Fig. S10). The 16S rRNA marker provided limited discrimination within the collection: all 60 isolates resolved into only 38 distinct 16S rRNA genotypes; 11 genotype groups contained two or more isolates and together accounted for 33 strains (55%), leaving 22 redundant tips (Fig. 5c). Metabolic niche, characterized independently as graded carbohydrate assimilation across eight carbon sources, was likewise unrelated to network position. None of the 18 strain-level associations between assimilation capacity and interaction descriptors reached significance (all *q* ≥ 0.73), and no layer showed a significant dyad-level association between metabolic distance and interaction presence (all *q* ≥ 0.085; Fig. S11, Supplementary Methods S9). At the resolution of these measurements, neither 16S rRNA relatedness nor carbohydrate assimilation showed a detectable association with interaction-network position.

**Figure 5.**
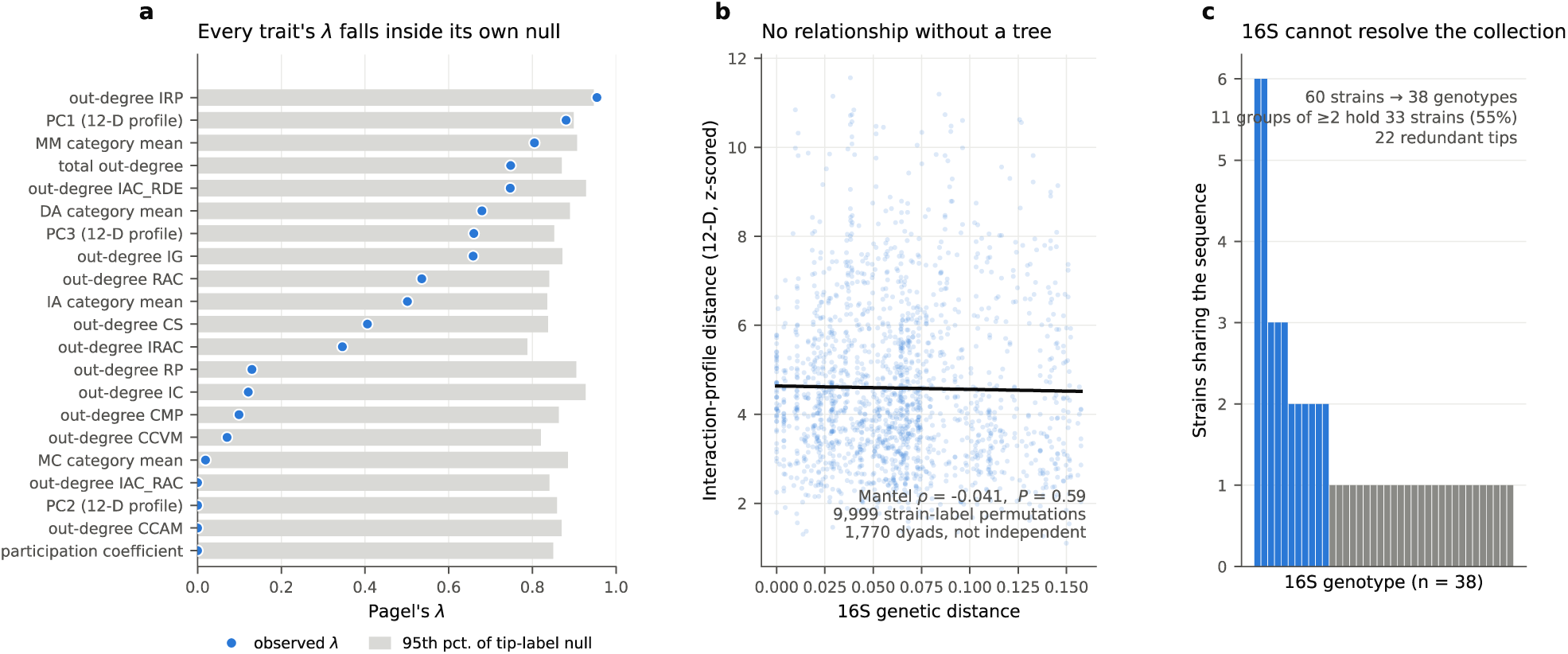
Interaction profiles show no detectable association with relatedness at 16S rRNA resolution. a, maximum-likelihood estimates of Pagel’s λ for 21 strain-level ecological traits on the 16S maximum-likelihood tree (points), each against the 95th percentile of its own tip-label permutation null (bars, 999 shuffles). Every observed λ falls inside its null (minimum *q*=0.39 after Benjamini-Hochberg correction). b, tree-independent Mantel test of 16S genetic distance against interaction-profile distance, the latter the Euclidean distance between *z*-standardized twelve-layer out-degree profiles (*ρ*=-0.041, P=0.59). The 1,770 plotted points are dyads, not independent observations, and the probability comes from 9,999 permutations of strain labels. c, distribution of 16S genotype sizes: the 60 isolates resolve into 38 genotypes, with 11 groups of two or more accounting for 33 strains (55%) and leaving 22 redundant tips, which bounds the power of any phylogenetic-signal test. Permutation inference was used because identical sequences make the phylogenetic covariance matrix rank deficient and the asymptotic χZ test anticonservative (Figure S9). These results apply at the resolution of the 16S marker and do not exclude structuring associated with finer-scale genomic variation.

Excluding the four non-*Streptomyces* isolates preserved the ordering of layer densities and largest strongly connected components (Spearman *ρ=*1.00 and 0.998) and retained two principal components under the 95th-percentile criterion. At matched 10,000 null draws, six enriched layer pairs remained compared with eight in the full collection, while one depleted pair remained in both (Table S4).

### 3.8. Spatial separation has phenotype-specific effects on interaction networks

We compared growth inhibition (IG) and change in sporulation (CS) between close-proximity and spatially separated inocula on the same plates (Fig. 6; Fig. S12; Supplementary Methods S10). These paired matrices were scored separately from the direct-contact layers of Table 1 and are analysed on their own. IG was detected in 785 of 3,540 directed non-self pairs under close proximity (density 0.222) and 261 under separation (density 0.074), a density difference of 0.148 (95% dyad-bootstrap interval 0.136-0.160; paired dyad randomization *P*<10^-4^). Of the 797 pairs positive in either configuration, 249 persisted, 536 were lost and 12 were gained after separation (loss:gain odds ratio 44.7, 95% CI 25.2-79.1).

**Figure 6.**
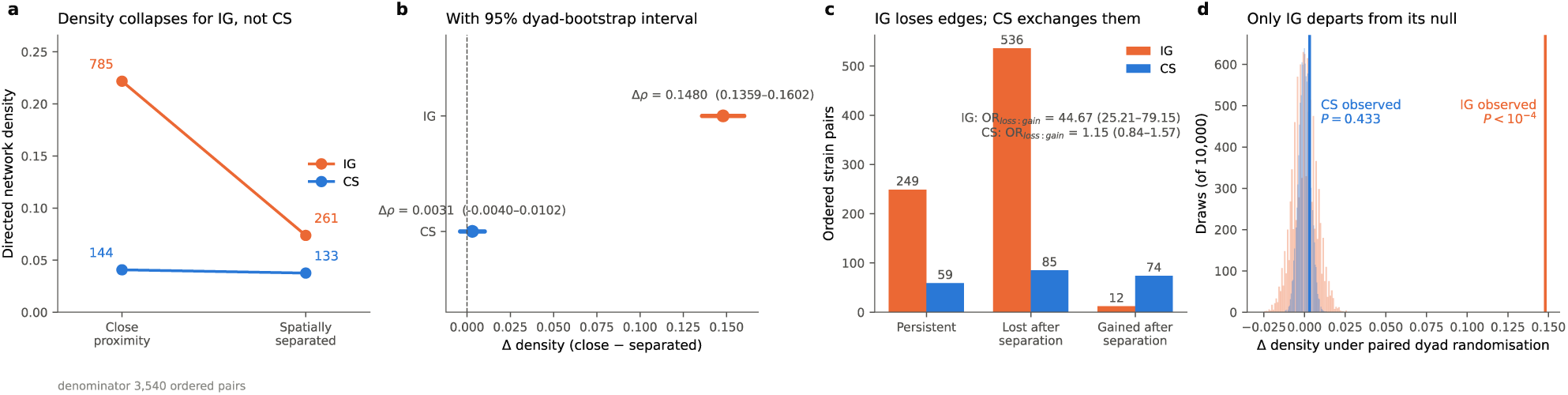
Spatial separation has phenotype-specific effects on interaction networks. Growth inhibition (IG) and change in sporulation (CS) were scored in paired configurations on each plate, with the inocula close enough for the colonies to meet and with the inocula separated (Supplementary Methods S10). a, directed network density in each configuration, labelled with the number of positive ordered pairs (denominator 3,540 ordered non-self pairs among 60 isolates). b, density difference Δρ (close - separated) with a 95% interval from 10,000 bootstrap resamples of unordered strain dyads. c, ordered pairs that persisted, were lost or were gained after separation, with the loss:gain odds ratio and its Wald 95% interval. d, null distribution of Δ*ρ* under 10,000 paired randomisations that exchange the configuration label of a whole dyad, with the observed value marked; IG *P* < 10^-4^, CS *P*=0.433. IG loses most of its edges under separation, whereas the aggregate frequency of CS barely changes although 85 responses are lost and 74 gained. The paired spatial matrices (Figure S12) were scored separately from the direct-contact layers of Table 1 and are analysed on their own. The comparison measures the effect of the tested separation and does not distinguish contact from diffusible or other distance-dependent mechanisms.

CS was detected in 144 directed pairs under close proximity (density 0.041) and 133 under separation (density 0.038); the density difference of 0.003 (95% interval -0.004 to 0.010) was compatible with the randomization null (*P=*0.433). Similar aggregate frequencies concealed substantial turnover. 59 pairs persisted, 85 were lost and 74 were gained (odds ratio 1.15, 95% CI 0.84-1.57). Separation therefore reduced the frequency of IG, whereas the aggregate frequency of CS changed little despite turnover in the responding pairs.

## 4. Discussion

This study supports a shift from viewing microbial competition as a one-dimensional hierarchy toward ecological identity as a multidimensional property emerging from partially independent processes. Classical competition theory predicts that strong niche overlap, when not counterbalanced by stabilizing niche differences, can allow persistent fitness differences to drive competitive exclusion. However, when ecological performance depends on several factors, competitive rankings become context dependent and multidimensional differentiation can promote diversity [39-41]. This perspective is particularly relevant to microbial communities, where competition encompasses resource exploitation, interference competition, antimicrobial production, contact-dependent interactions, and other context-dependent mechanisms [42]. Negative interactions indeed dominate many microbial communities [43]. Our results extend this framework by showing that dimensionality may arise not only through competition for multiple resources, but also through different phenotypic responses of microbial interaction. Thus, competitive ability alone does not adequately define ecological identity.

Multilayer analysis consequently reveals information lost when microbial interactions are represented exclusively as antagonistic networks. Ecological communities simultaneously contain multiple forms of interaction [44], including metabolic, physical, regulatory, and signalling processes that can affect community organization through different mechanisms [45]. Most cross-layer overlaps are indistinguishable from degree-constrained expectations, while a restricted subset show excess or depleted correspondence. However, phenotype-specific deviations in reciprocity and transitivity demonstrated distinct network organization, potentially affecting ecological stability differently across interaction phenotypes [46]. The pronounced excess reciprocity of IG therefore provides stronger evidence of non-random organization than its large SCC alone.

At the strain level, results converged on heterogeneous, multidimensional ecological strategies. Parallel analysis supported two principal axes robustly and a third only marginally, while Gaussian-mixture modelling produced no stable component count. Rankings varied substantially among phenotypes, indicating phenotype-dependent ecological influence, while agreement between normalized David’s scores and Bradley-Terry estimates shows the two estimators to be concordant, though both are computed from the same matrices and so do not validate each other independently [34]. Competitive outcomes can themselves depend on ecological context, for example, antibiotic-mediated interactions among *Streptomyces* can generate bi-stability, whereby established strains resist invasion despite being unable to invade from low abundance [47]. Ecological superiority is therefore not necessarily intrinsic, and the inducible-antagonism layers indicate that strategy can itself be interaction dependent. Neighbouring *Streptomyces* may induce antibiotic production while suppressing antibiosis in competitors, making realized antagonistic capacity dependent on social context [48]. Such responses can convert lethal relationships into neutral ones and potentially facilitate coexistence. This resembles trait-mediated interactions, in which one organism induces physiological, morphological, or behavioural changes that modify subsequent interactions [49].

Although the distribution of interaction profiles is compatible with continuous ecological variation, our analyses do not establish that the underlying population is strictly continuous or unimodal. The information criteria and the cross-validated likelihood disagree, and the best-scoring model overall was a diagonal three-component mixture. This may reflect either genuinely weak cluster structure or limited power to estimate complex mixtures from 60 strains. We therefore interpret these analyses as showing no robust evidence for discrete ecological groups at the present sample size, rather than demonstrating their absence. Larger strain collections will be required to determine whether finer-scale or partially overlapping ecological groups can be resolved.

We hypothesize that the observed multidimensionality could reflect semi-independent evolution and conditional coupling of secondary metabolism, development, sensing, and stress responses. Specialized-metabolite systems diversify through gene gain, loss, recombination, and horizontal transfer [50, 51], while developmental, signalling, and stress pathways modulate antimicrobial activity and physiology [48, 50, 51].

An important limitation is that some interaction layers were not experimentally independent. Developmental and morphological phenotypes were scored from common confrontation assays, and several antimicrobial phenotypes shared related overlay-based procedures. Consequently, part of the observed within-group correspondence could arise from shared assay conditions or correlated scoring variation. Degree-preserving network randomization controls for expected overlap arising from network topology, but cannot correct for such measurement covariance. Accordingly, we interpret enriched overlap as evidence that particular phenotypic outcomes co-occur more frequently than expected from their network structure, without assigning this correspondence exclusively to shared molecular regulation. Future experiments using independent quantitative readouts will be required to partition biological coupling from assay-associated covariance. A second limitation concerns what a directed edge represents: RAC is dominated by the strain in which it is scored, so its layer-level statistics summarize a release phenotype rather than partner-specific structure, a constraint that does not extend to the genuinely dyadic layers.

This supports a modular-but-interconnected architecture rather than strict independence. Local interactions, microscale organization, soil architecture and spatial scale are known to influence diversity and interaction networks [52-55]. Spatial dependence differed between the two phenotypes tested. Growth inhibition was less frequent under separation, whereas the aggregate frequency of sporulation responses changed little despite substantial turnover in the responding pairs. These results demonstrate different responses to the tested spatial configuration, but they do not distinguish physical contact from diffusible or other distance-dependent effects.

These results bear on coexistence. Multidimensional competition can weaken universal competitive rankings and increase opportunities for coexistence, particularly when competitive dimensions are weakly correlated [41]. Antibiotic production, sensitivity, and degradation can generate interaction networks compatible with stable coexistence [9]. Our results suggest an additional mechanism of interaction-niche dimensionality where strains disadvantaged in growth inhibition may remain influential through developmental or inducible phenotypes, whereas strong antagonists may not dominate other dimensions. Although our experiments do not directly demonstrate stable coexistence, they reveal the context-dependent heterogeneity upon which coexistence mechanisms could operate.

We detected no association between interaction phenotypes and relatedness at 16S rRNA resolution, for Pagel’s λ, Mantel and layer-specific analyses alike. This is not evidence of evolutionary independence: the limited sequence variation, including identical 16S rRNA sequences among strains, restricts resolution of recent divergence. Genome-resolved analyses would be required to test whether ecological similarity tracks finer-scale relatedness. The same applied to metabolic niche as captured by carbohydrate assimilation, which showed no detectable association with either strain-level network position or dyad-level interaction presence. Both results are bounded by what the two markers resolve, and neither excludes structuring by properties they do not capture.

Several limitations delimit these conclusions. The networks represent standardized pairwise interaction potential rather than realized interactions in heterogeneous soils, where multiple partners, resource fluctuations, priority effects, and higher-order interactions coexist. Binary scoring also removes information on interaction strength, concentration dependence, and temporal dynamics. Moreover, the twelve layers represent phenotypic outcomes rather than demonstrated molecular mechanisms; the proposed modular evolutionary model therefore requires genomic, transcriptomic, metabolomic, and functional validation. Spatial comparisons were restricted to IG and CS, preventing generalization across all phenotypes. Finally, pairwise interactions may not predict multispecies dynamics because indirect and higher-order effects can violate pairwise additivity [49, 56].

## 5. Conclusion

Direct-contact ecological behaviour is better represented as a multidimensional network of partially coupled interaction axes than as a single antagonistic network. This matters for microbiome engineering: screening based predominantly on pathogen inhibition may select strong individual antagonists while overlooking incompatibilities or complementary functions expressed through developmental, metabolic, inducible or signalling interactions. Multidimensional profiling could guide the selection of strain combinations with complementary interaction properties; whether this improves consortium establishment, pathogen suppression, or rhizosphere stability requires direct testing.

## References

1. Bengtsson-Palme J. Microbial model communities: to understand complexity, harness the power of simplicity. Comput Struct Biotechnol J 2020;18:3987–4001. 10.1016/j.csbj.2020.11.043

2. Darwin C. On the Origin of Species by Means of Natural Selection. London: John Murray, 1859. Chapter III, Struggle for existence.

3. Godwin CM, Chang FH, Cardinale BJ. An empiricist’s guide to modern coexistence theory for competitive communities. Oikos 2020;129:1109–27. 10.1111/oik.06957

4. Chen X, Crocker K, Kuehn S et al. Inferring resource competition in microbial communities from time series. PRX Life 2025;3:023019. 10.1103/8gkj-rdzy

5. Goyal A, Maslov S. Diversity, stability, and reproducibility in stochastically assembled microbial ecosystems. Phys Rev Lett 2018;120:158102. 10.1103/PhysRevLett.120.158102

6. Chater KF. Recent advances in understanding Streptomyces. F1000Res 2016;5:2795. 10.12688/f1000research.9534.1

7. Verschoor JA, Croese MRJ, Lakemeier SE et al. Polyester degradation by soil bacteria: identification of conserved BHETase enzymes in Streptomyces. Commun Biol 2024;7:725. 10.1038/s42003-024-06414-z

8. Czárán TL, Hoekstra RF, Pagie L. Chemical warfare between microbes promotes biodiversity. Proc Natl Acad Sci U S A 2002;99:786–90. 10.1073/pnas.012399899

9. Kelsic ED, Zhao J, Vetsigian K et al. Counteraction of antibiotic production and degradation stabilizes microbial communities. Nature 2015;521:516–19. 10.1038/nature14485

10. Mohite OS, Jørgensen TS, Booth TJ et al. Pangenome mining of the Streptomyces genus redefines species’ biosynthetic potential. Genome Biol 2025;26:9. 10.1186/s13059-024-03471-9

11. Schlatter DC, Kinkel LL. Do tradeoffs structure antibiotic inhibition, resistance, and resource use among soil-borne Streptomyces? BMC Evol Biol 2015;15:186. 10.1186/s12862-015-0470-6

12. Vetsigian K, Jajoo R, Kishony R. Structure and evolution of Streptomyces interaction networks in soil and in silico. PLoS Biol 2011;9:e1001184. 10.1371/journal.pbio.1001184

13. Sun T, Liu H, Wang N et al. Interactions with native microbial keystone taxa enhance the biocontrol efficiency of Streptomyces. Microbiome 2025;13:126. 10.1186/s40168-025-02120-y

14. Westhoff S, Kloosterman AM, van Hoesel SFA et al. Competition sensing changes antibiotic production in Streptomyces. mBio 2021;12:e02729–20. 10.1128/mBio.02729-20

15. Niu G, Chater KF, Tian Y et al. Specialised metabolites regulating antibiotic biosynthesis in Streptomyces spp. FEMS Microbiol Rev 2016;40:554–73. 10.1093/femsre/fuw012

16. D’Souza G, Shitut S, Preussger D et al. Ecology and evolution of metabolic cross-feeding interactions in bacteria. Nat Prod Rep 2018;35:455–88. 10.1039/C8NP00009C

17. Katoh K, Standley DM. MAFFT multiple sequence alignment software version 7: improvements in performance and usability. Mol Biol Evol 2013;30:772–80. 10.1093/molbev/mst010

18. Capella-Gutiérrez S, Silla-Martínez JM, Gabaldón T. trimAl: a tool for automated alignment trimming in large-scale phylogenetic analyses. Bioinformatics 2009;25:1972–3. 10.1093/bioinformatics/btp348

19. Price MN, Dehal PS, Arkin AP. FastTree 2 - approximately maximum-likelihood trees for large alignments. PLoS ONE 2010;5:e9490. 10.1371/journal.pone.0009490

20. Piñeiro C, Abuín JM, Pichel JC. Very Fast Tree: speeding up the estimation of phylogenies for large alignments through parallelization and vectorization strategies. Bioinformatics 2020;36:4658–9. 10.1093/bioinformatics/btaa582

21. Davelos AL, Kinkel LL, Samac DA. Spatial variation in frequency and intensity of antibiotic interactions among streptomycetes from prairie soil. Appl Environ Microbiol 2004;70:1051–8. 10.1128/AEM.70.2.1051-1058.2004

22. Guimerà R, Amaral LAN. Functional cartography of complex metabolic networks. Nature 2005;433:895–900. 10.1038/nature03288

23. Jaccard P. The distribution of the flora in the alpine zone. New Phytol 1912;11:37–50. 10.1111/j.1469-8137.1912.tb05611.x

24. Maslov S, Sneppen K. Specificity and stability in topology of protein networks. Science 2002;296:910–13. 10.1126/science.1065103

25. Phipson B, Smyth GK. Permutation P-values should never be zero: calculating exact P-values when permutations are randomly drawn. Stat Appl Genet Mol Biol 2010;9:Article39. 10.2202/1544-6115.1585

26. Benjamini Y, Yekutieli D. The control of the false discovery rate in multiple testing under dependency. Ann Stat 2001;29:1165–88. 10.1214/aos/1013699998

27. Horn JL. A rationale and test for the number of factors in factor analysis. Psychometrika 1965;30:179–85. 10.1007/BF02289447

28. Peres-Neto PR, Jackson DA, Somers KM. How many principal components? Stopping rules for determining the number of non-trivial axes revisited. Comput Stat Data Anal 2005;49:974–97. 10.1016/j.csda.2004.06.015

29. Smyth P. Model selection for probabilistic clustering using cross-validated likelihood. Stat Comput 2000;10:63–72. 10.1023/A:1008940618127

30. Miloslavsky M, van der Laan MJ. Fitting of mixtures with unspecified number of components using cross validation distance estimate. Comput Stat Data Anal 2003;41:413–28. 10.1016/S0167-9473(02)00166-4

31. Celeux G, Govaert G. Gaussian parsimonious clustering models. Pattern Recognit 1995;28:781–93. 10.1016/0031-3203(94)00125-6

32. Hartigan JA, Hartigan PM. The dip test of unimodality. Ann Stat 1985;13:70–84. 10.1214/aos/1176346577

33. David HA. Ranking from unbalanced paired-comparison data. Biometrika 1987;74:432–6. 10.1093/biomet/74.2.432

34. de Vries H, Stevens JMG, Vervaecke H. Measuring and testing the steepness of dominance hierarchies. Anim Behav 2006;71:585–92. 10.1016/j.anbehav.2005.05.015

35. Bradley RA, Terry ME. Rank analysis of incomplete block designs: I. The method of paired comparisons. Biometrika 1952;39:324–45. 10.1093/biomet/39.3-4.324

36. Pagel M. Inferring the historical patterns of biological evolution. Nature 1999;401:877–84. 10.1038/44766

37. Benjamini Y, Hochberg Y. Controlling the false discovery rate: a practical and powerful approach to multiple testing. J R Stat Soc Series B Stat Methodol 1995;57:289–300. 10.1111/j.2517-6161.1995.tb02031.x

38. Mantel N. The detection of disease clustering and a generalized regression approach. Cancer Res 1967;27:209–20.

39. Chesson P. Mechanisms of maintenance of species diversity. Annu Rev Ecol Evol Syst 2000;31:343–66. 10.1146/annurev.ecolsys.31.1.343

40. Mayfield MM, Levine JM. Opposing effects of competitive exclusion on the phylogenetic structure of communities. Ecol Lett 2010;13:1085–93. 10.1111/j.1461-0248.2010.01509.x

41. Allesina S, Levine JM. A competitive network theory of species diversity. Proc Natl Acad Sci U S A 2011;108:5638–42. 10.1073/pnas.1014428108

42. Stubbendieck RM, Straight PD. Multifaceted interfaces of bacterial competition. J Bacteriol 2016;198:2145–55. 10.1128/JB.00275-16

43. Foster KR, Bell T. Competition, not cooperation, dominates interactions among culturable microbial species. Curr Biol 2012;22:1845–50. 10.1016/j.cub.2012.08.005

44. Carrara F, Giometto A, Seymour M et al. Inferring species interactions in ecological communities: a comparison of methods at different levels of complexity. Methods Ecol Evol 2015;6:895–906. 10.1111/2041-210X.12363

45. Widder S, Allen RJ, Pfeiffer T et al. Challenges in microbial ecology: building predictive understanding of community function and dynamics. ISME J 2016;10:2557–68. 10.1038/ismej.2016.45

46. de Vos MGJ, Zagorski M, McNally A et al. Interaction networks, ecological stability, and collective antibiotic tolerance in polymicrobial infections. Proc Natl Acad Sci U S A 2017;114:10666–71. 10.1073/pnas.1713372114

47. Wright ES, Vetsigian KH. Inhibitory interactions promote frequent bistability among competing bacteria. Nat Commun 2016;7:11274. 10.1038/ncomms11274

48. Abrudan MI, Smakman F, Grimbergen AJ et al. Socially mediated induction and suppression of antibiosis during bacterial coexistence. Proc Natl Acad Sci U S A 2015;112:11054–9. 10.1073/pnas.1504076112

49. Utsumi S, Kishida O, Ohgushi T. Trait-mediated indirect interactions in ecological communities. Popul Ecol 2010;52:457–9. 10.1007/s10144-010-0236-3

50. Hibbing ME, Fuqua C, Parsek MR et al. Bacterial competition: surviving and thriving in the microbial jungle. Nat Rev Microbiol 2010;8:15–25. 10.1038/nrmicro2259

51. Granato ET, Meiller-Legrand TA, Foster KR. The evolution and ecology of bacterial warfare. Curr Biol 2019;29:R521–R537. 10.1016/j.cub.2019.04.024

52. Kerr B, Riley MA, Feldman MW et al. Local dispersal promotes biodiversity in a real-life game of rock-paper-scissors. Nature 2002;418:171–4. 10.1038/nature00823

53. Cordero OX, Datta MS. Microbial interactions and community assembly at microscales. Curr Opin Microbiol 2016;31:227–34. 10.1016/j.mib.2016.03.015

54. Young IM, Crawford JW. Interactions and self-organization in the soil-microbe complex. Science 2004;304:1634–7. 10.1126/science.1097394

55. Galiana N, Lurgi M, Claramunt-López B et al. The spatial scaling of species interaction networks. Nat Ecol Evol 2018;2:782–90. 10.1038/s41559-018-0517-3

56. Momeni B, Xie L, Shou W. Lotka-Volterra pairwise modeling fails to capture diverse pairwise microbial interactions. eLife 2017;6:e25051. 10.7554/eLife.25051

