## Supplementary data for "Multilayer interaction networks reveal multidimensional ecological strategies and context-dependent competition in soil actinobacteria"

### Supplementary Methods S1. Isolation, cultivation and molecular characterization

Actinobacteria (56 *Streptomyces*, plus one *Saccharothrix*, two *Lentzea* and one *Amycolatopsis* isolate) were isolated from soils collected in the Rif and Atlas mountain regions of Morocco, including the Middle Atlas, Anti-Atlas and High Atlas. Approximately 0.5 g of surface soil was suspended in 100 mL enrichment medium containing, per liter, 0.5 g $KH_2PO_4$, 1.2 g $K_2HPO_4$, 10 g peptone and 5 g yeast extract. Serial dilutions were plated on Bennett's agar containing, per liter, 1.0 g beef extract, 10.0 g glucose, 2.0 g casein hydrolysate, 1.0 g yeast extract, 15.0 g agar and 0.02 g calcium carbonate.

For molecular characterization, pure colonies were cultivated in Bennett broth without glucose for 5 days. Genomic DNA was extracted using the Monarch® Genomic DNA Purification Kit according to the manufacturer's instructions. Near-full-length 16S rRNA genes were amplified using OneTaq® polymerase and universal primers 27F (AGAGTTTGATCMTGGCTCAG) and 1492R (GGTTACCTTGTTACGACTT). Amplicons were sequenced by Sanger sequencing using BigDye™ Terminator v3.1 chemistry on a SeqStudio instrument (Thermo Fisher Scientific). Sequences were aligned with MAFFT (FFT-NS-i, 1,000 iterations) [1], trimmed with trimAl [2] (gappyout; 1,533 to 1,337 columns), and a maximum-likelihood tree was inferred under GTR+GAMMA with VeryFastTree, a FastTree 2 implementation [3, 4]. API 50CH, API 20 NE and API ZYM strips were used according to the manufacturer's instructions for biochemical characterization. Strain identities, 16S sequence lengths and GenBank accessions are listed in Table S1.

### Supplementary Methods S2. Pairwise interaction assay and phenotype definitions

Pairwise assays were conducted on Bennett's agar containing, per liter, 5 g beef extract, 5 g glucose, 2 g casein hydrolysate, 1 g yeast extract and 18 g agar, adjusted to pH 7.2.

For each ordered strain pair ($S_i$,$S_j$), colonies were evaluated under direct-contact conditions. Twelve phenotypic responses were scored: CMP, CS, CCVM, CCAM, RP, IRP, IG, IC, RAC, IAC_RAC, IRAC and IAC_RDE. Except where noted below, each response was scored on the colony of Sj in the confrontation against that same strain's monoculture control on the same plate, so a positive entry records a change in Sj attributable to the presence of Si. RP was scored as the appearance or intensification of pigment release by Sj relative to its monoculture control, and IRP as its suppression; both are therefore directed effects on the receiving strain rather than statements about intrinsic pigment production. IAC_RAC records that Sj released antibacterial compounds in the confrontation that were not detectable from its monoculture, IRAC that release detectable from monoculture was suppressed in the confrontation, and IAC_RDE that Sj acquired detectable antimicrobial-degrading activity. RAC differs from all of these in what it measures. It records antibacterial activity detected against the indicator strain in the overlay, and the assembled matrix behaves accordingly: the identity of the strain in which it is scored ($S_j$) accounts for 57.6% of its deviance against 0.8% for its partner ($S_i$), and once the scored strain is accounted for the partner explains only a further 1.9% (likelihood-ratio $\chi^2 = 96.1$ on 59 d.f., P = 0.002); each percentage is the share of the binomial deviance removed by a one-factor model of that strain relative to a constant-probability model. A RAC entry therefore records principally whether $S_j$ releases antibacterial compounds at all, rather than a partner-specific response. RAC is retained as a layer throughout, and this constraint is stated wherever its layer-level statistics are reported.

For antimicrobial assays, plates were overlaid with 5 mL soft LB agar containing approximately $2*10^4$ CFU.mL$^{-1}$ *B. subtilis* ATCC 6051 and incubated overnight at 30 °C. Inhibition zones were measured according to the assay described by Davelos et al. [5].

Directionality was determined by independently evaluating the phenotypic response of each strain in the pairwise confrontation relative to its corresponding monoculture control. Pairwise interactions were evaluated on Bennett's agar for all unique strain dyads. For ($N$=60) strains, this corresponded to 1,770 unordered pairwise confrontation assays in each run $\left(\left[N\left(N-1\right)/2\right]\right)$. Each confrontation plate contained both strains $\left(S_i\right)$ and $\left(S_j\right)$, and the phenotypic response of each colony was evaluated independently relative to its corresponding monoculture control. Thus, a single physical confrontation could yield two directed observations, $\left(S_i \rightarrow S_j\right)$ and $\left(S_j \rightarrow S_i\right)$, generating up to 3,540 directed non-self observations per phenotype layer. The unordered strain dyad therefore constituted the experimental confrontation unit, whereas the directed strain-response pair constituted the analytical unit used to construct the multilayer adjacency matrices.

The complete set of 1,770 confrontations was performed in two independent runs. Every plate was read twice by the same investigator and the scores were reviewed by a second investigator. The two runs and the two readings agreed except for rare discordant calls; each discordant plate was re-read until a single call was reached, and the final binary matrices record that consensus. The number of discordant calls was not recorded, so the data carry no numerical estimate of scoring reproducibility. Sources of variation such as plate-to-plate variation, inoculum preparation and reader judgement at the threshold of a call were therefore resolved by consensus rather than estimated. Because every layer was produced by the same procedure, contrasts between layers are not obviously biased by this, but the exact edge count of any one layer, and any individual edge, should be read with this in mind.

### Supplementary Methods S3. Binary multilayer and tensor representations

For each of the twelve interaction responses r (1…12), we constructed a directed binary adjacency matrix $M^{(r)}$. The scoring rules were : *(i)* if $M_{ij}^{(r)}=1$, it does not imply $M_{ji}^{(r)}=1$, and *(ii)* $M_{ij}^{(r)}=1$ if strain $S_j$ elicited response $r$ in strain $S_i$ (i.e., a detectable effect of $S_j$ on $S_i$), and 0 otherwise. All per-layer analyses (density, reciprocity, SCC, Jaccard overlap, PCA, GMM, dominance scoring) operate on the binary projection $A_l=1[M_l>0]$; under standardization, similarity, and thresholding the layer constant cancels, so the analyses are intrinsically binary by design.

A phenotype was recorded as positive when the two runs and the two readings agreed that it was present or, for the rare discordant cases, when re-reading resolved it as present (Supplementary Methods S2).

The twelve layers are organized into four functional categories used by the downstream embedding: direct antagonism (DA) = {IG, IC, RAC}; induced antagonism (IA) = {IAC_RAC, IAC_RDE, IRAC}; metabolic modulation (MM) = {release of pigment (RP), inhibition of pigment release (IRP)}; morphogenetic control (MC) = {colour of aerial mycelium (CCAM), colour of vegetative mycelium (CCVM), change in mycelium production (CMP), change in sporulation (CS)}.

$$A_{supra}=\begin{pmatrix} M^{(1)} & \omega I & . & \omega I \\ \omega I & M^{(2)} & . & \omega I \\ . & . & . & . \\ \omega I & \omega I & . & M^{L} \end{pmatrix}$$

Interaction matrices are represented as layers of a single three dimensional tensor. The tensor *M* is defined by $M(i,S_i,S_j) = M_i(S_i,S_j)$. Binary matrices were then stacked into a tensor with entries $M^{(bin)}((i,S_i,S_j) \in (1,0))$. This tensor format is the basis for all subsequent multilayer network analyses. Matrices were re-indexed to match a reference node ordering. Binary tensors, $W \in \mathbb{R}^{Lxnxn}$, of shape (*L, n, n* ) were constructed, where *L* is the number of layers and *N* the number of nodes in each layer.

### Supplementary Methods S4. Network topology, cross-layer similarity and null models

For each layer's binary projection $A_l=1[M_l>0]$ we computed edge density $\rho_l=E_l \vee (N(N-1))$, mutual-pair count (unordered dyads {i,j} with $A_l[i,j]=A_l[j,i]=1$, and reciprocity, defined on a pairs basis as $R_p=mutual\,pairs/connected\,pairs$, i.e. the fraction of connected dyads that are mutual. We report pairs-based reciprocity throughout. An edges-based variant, $R_e=2R_p/(1+R_p)$, yields identical layer orderings but uniformly larger values; it is not used here. Strongly connected components were identified with Tarjan's algorithm as implemented in NetworkX [6]. The out-degree of node i in layer l is $k_{i,l}^{out}=\Sigma_{j\neq i}A_l[i,j]$. The participation coefficient [7], $P_i=1-\sum(\frac{k_{i,l}^{out}}{k_i^{out}})^2$, was computed over the twelve layers, where $k_{i,l}^{out}$ is the out-degree of strain i in layer l and $k_i^{out}=\sum k_{i,l}^{out}$ is its total out-degree summed across layers. $P_i$ ranges from 0 (all interactions confined to a single layer) to near 1 (interactions spread evenly across layers).

For each layer pair ($l_i$, $l_j$), observed Jaccard similarity [8] was computed on the directed edge sets as $J_{ij}^{obs}=E_i\cap E_j/E_i\cup E_j$. Cosine similarity and Pearson correlation between flattened matrices were also computed alongside, for layer-similarity visualization.

A degree-preserving null model was used for layer overlap, SCC size, reciprocity and transitivity. For each layer we generated 10,000 null networks by directed edge swaps [9]; each null network was obtained from the observed layer by E successful directed three-edge swaps, where E is the number of edges in that layer (NetworkX directed_edge_swap), which preserves every strain's in- and out-degree exactly. For each null draw we computed $J^{null}$ from independently-rewired versions of the two layers. Two-sided empirical P-values were obtained with the Phipson-Smyth $(M+1)/(N+1)$ correction [10], with b the number of draws at least as far from the null mean as the observed value, and then FDR-corrected across the 66 layer pairs (66 total pairwise comparisons; the 65 remaining after separately flagging CCAM-CCVM form the set used for the main biological summary, which is not a claim that those 65 tests are statistically independent) using Benjamini-Yekutieli [11], which is robust to the dependence structure imposed by each layer appearing in 11 of the 66 tests. Benjamini-Hochberg [12] results are reported in Supplementary Table S2 for completeness. The same 10,000 null draws were reused for testing the largest SCC, pair-based reciprocity and transitivity of each layer (Figure S3; Table 1).

Because raw Jaccard similarity is intrinsically influenced by network density, raw coefficients were treated as descriptive measures of edge-set overlap rather than as evidence of statistical independence between layers. For each layer pair (l, m), departure from degree-constrained expectation was quantified as a standardized effect size, $\left(Z_{lm}=\left[J_{lm}^{obs}-E\left(J_{lm}^{null}\right)\right]/SD\left(J_{lm}^{null}\right)\right)$, where positive values indicate enriched overlap and negative values indicate depleted overlap relative to the degree-preserving null. Statistical inference was based on empirical permutation P-values corrected using the Benjamini-Yekutieli procedure. Non-significant deviations from the null expectation were interpreted as null-compatible correspondence and not as evidence that layers were statistically or biologically independent.

The CCAM-CCVM comparison was designated *a priori* as potentially affected by shared measurement because both color phenotypes were scored simultaneously from the same colonies and was therefore interpreted separately.

It is worth noting that the pair {CCAM, CCVM} was flagged *a priori* as a methodological-confound pair. Aerial and vegetative mycelium colour were scored from the same plate at each reading by the same observer, so any shared signal between them confounds shared biology with shared measurement. We report this pair separately from the biological summary and discuss it explicitly in Results. More generally, several phenotypes shared assay contexts beyond the prespecified CCAM-CCVM pair. CMP, CS, CCAM and CCVM were derived from common colony-confrontation assays, whereas RAC, IAC_RAC, IRAC and IAC_RDE involved related antimicrobial overlay procedures. Degree-preserving null models account for edge density and degree heterogeneity but do not model covariance introduced by common experimental procedures. Accordingly, overlap among same-assay phenotypes was interpreted as phenotypic correspondence potentially comprising both biological and measurement-associated components. Cross-assay layer pairs provide a complementary assessment less directly susceptible to this source of covariance.

Sensitivity to taxonomic composition was assessed by repeating the headline analyses on the 56 *Streptomyces* isolates alone, with 10,000 fresh degree-preserving null draws generated for the reduced network so that both cohorts are compared at a matched draw count (Table S4).

**Supplementary Methods S5. Ecological embedding**

For each strain $S_i$, we summarize its ecological interaction profile using four functional categories: DA, IA, MM, and MC, defined exactly as in the multilayer representation above: direct antagonism (DA = {IG, IC, RAC}), induced antagonism (IA = {IAC_RAC, IAC_RDE, IRAC}), metabolic/pigment modulation (MM = {RP, IRP}), and morphogenetic/developmental control (MC = {CCAM, CCVM, CMP, CS}). For every strain we compute the mean outgoing interaction intensity within each ecological category using the following formula, $e_{i,k}=\frac{1}{|C_k|}\sum_{r\in C_k}\left(\frac{1}{59}\sum_{j\neq i} M^{(r)}_{i\to j}\right)$, where $M^{(r)}_{i\to j}$ is the binary entry of layer *r*. This yields a 4-dimensional ecological embedding vector $E_i=[e_i, DA, e_i, IA, e_i, MM, e_i, MC]$. For descriptive visualization of broader functional patterns, the 12 interaction phenotypes were grouped a priori into four functional categories: direct antagonism (DA: IG, IC, RAC), induced antagonism (IA: IAC_RAC, IAC_RDE, IRAC), metabolic modulation (MM: RP, IRP), and morphological/developmental change (MC: CCAM, CCVM, CMP, CS). Because categories contained unequal numbers of phenotypes (DA = 3, IA = 3, MM = 2, MC = 4), category scores were calculated as the mean outgoing interaction frequency across constituent layers, preventing categories with more phenotypes from receiving greater numerical weight. This four-dimensional representation was used as a descriptive visualization of functional interaction profiles and was not treated as an independent validation of the structure identified from the complete 12-dimensional dataset.

**Supplementary Methods S6. PCA and assessment of cluster structure**

PCA was conducted with scikit-learn [13] on z-standardized 12-dimensional strain profiles. Horn's parallel analysis [14] was based on 1,000 datasets generated by independently permuting values within each column, which preserves every layer's marginal distribution while destroying between-layer covariance. A component was retained when its observed variance exceeded the 95th percentile of the corresponding null distribution; the weaker mean-eigenvalue criterion is reported alongside as a sensitivity analysis, and the two disagree for PC3 (observed 11.82%, null mean 11.65%, null 95th percentile 12.62%; Figure S4). Uncertainty on the observed variances is from 2,000 nonparametric bootstrap resamples of the 60 strains, each re-standardized and re-decomposed, summarized as 2.5th and 97.5th percentiles.

Gaussian mixture models containing ($k$=1-8) components and full, diagonal or spherical covariance structures [15] were fitted to both 12-dimensional interaction profiles and four-dimensional ecological embeddings. Model order was determined principally from 10-fold cross-validated held-out log-likelihood [16] because conventional in-sample information criteria showed evidence of overfitting given (N=60) and the dimensionality of the data. BIC, AIC and ICL were retained as supplementary diagnostics; across the two embeddings and three covariance structures their preferred component count ranged from four to eight, so they do not identify a single solution either. Each fit used 50 random initializations, a covariance regularization of 1e-4 and a shared 10-fold partition, so that all component counts are scored on identical splits. Free parameters grow steeply with k under full covariance — 90 free parameters per component in the 12-dimensional profile (78 covariance terms and 12 means) against 24 under diagonal and 13 under spherical — and the held-out likelihood of the full-covariance family collapses accordingly, from -14.53 at k = 1 to -3673 at k = 8. Those full-covariance solutions at k > 3 are reported as diagnostics of overfitting rather than as candidate models. The highest held-out log-likelihood obtained by any model was the diagonal three-component solution (-12.19), against -14.53 for the best full-covariance model and -15.97 for the best spherical one (Figure S5).

Hartigan's dip test [17] independently tested unimodality along leading PCA dimensions, while silhouette coefficients quantified separation where multi-component solutions were considered.

**Supplementary Methods S7. Dominance and recurrent interactions**

Normalized David's score was calculated independently for each layer from dyadic interaction proportions [18]. High values characterize strains that repeatedly act upon multiple partners while being comparatively less affected themselves. To summarize asymmetry in directed interaction networks, we calculated David’s score (DS) for each strain within each phenotype layer [19].

David's score was originally developed for dominance matrices based on wins and losses; here, its mathematical formulation was used more generally to summarize the relative position of strains in directed interaction networks. Because an edge $(S_i \rightarrow S_j)$ denotes that strain $(S_i)$ elicited a scored phenotypic response in $(S_j)$, rather than necessarily representing a competitive victory, DS was interpreted as a sender influence score. Higher values therefore indicate strains that more frequently elicited directional responses relative to the responses they received, directly or indirectly through the network structure. Competitive interpretations were restricted to explicitly antagonistic phenotypes, particularly growth inhibition; RAC was excluded from this interpretation because its entries are dominated by the strain in which it is scored (Supplementary Methods S2). For developmental, morphological, pigmentation-associated, metabolic, or induced-response layers, DS was interpreted exclusively as relative interaction influence. Bradley-Terry models [20] were used as a complementary ranking approach based on the same directed matrices and were interpreted under the same phenotype-specific constraints.

Two multilayer descriptors summarize how stable a strain's direction of influence is across phenotypes. Within each layer a strain was classed as a net sender when its out-degree exceeded its in-degree and as a net receiver in the converse case, layers in which the two were equal being left unclassified. Role instability is the number of classified layers whose class differs from that strain's majority class, from 0 for one role throughout to 6 for an even split; variability of interaction output is the standard deviation of a strain's out-degree across the twelve layers (Figure 4). Within-layer agreement between David's scores and Bradley-Terry strengths is given in Table S3.

**Supplementary Methods S8. Phylogenetic signal**

Near-full-length 16S rRNA sequences (≥1,200 bp) were aligned, trimmed and used to reconstruct a maximum-likelihood phylogeny under GTR+GAMMA as described in Supplementary Methods S1. Pagel's $\lambda$ was estimated [21] for 21 traits: *(i)* 12 layer-specific out-degrees, *(ii)* four functional-category scores, *(iii)* total out-degree, *(iv) participation coefficient and (v) PC1, PC2 and PC3*; PC3 is included as a descriptive axis although it did not meet the 95th-percentile retention criterion (Supplementary Methods S6). The 60 isolates resolve into 38 distinct 16S genotypes: 11 genotype groups contain two or more isolates and together account for 33 strains (55%), leaving 22 redundant tips. This produced a rank-deficient phylogenetic covariance matrix, while the maximum-likelihood tree was strongly non-ultrametric. Diagnostic randomization showed that conventional asymptotic likelihood-ratio testing generated anticonservative estimates under these conditions (Figure S9). Significance was therefore determined empirically using 999 tip-label permutations, which preserved tree structure and each trait's marginal distribution while destroying trait-tip correspondence. Resulting probabilities were corrected across traits using Benjamini-Hochberg adjustment. Robustness was assessed using UPGMA and WPGMA trees derived from pairwise 16S rRNA distances. A complementary tree-independent Mantel test [22] with 9,999 permutations compared pairwise 16S rRNA genetic distances with distances among 12-dimensional interaction profiles, the latter taken as the Euclidean distance between z-standardized out-degree profiles. Permutation was applied to strain labels because the 1,770 dyadic distances are not independent observations.

**Supplementary Methods S9. Metabolic niche and network position**

Carbohydrate assimilation was scored from API 50CH strips for all 60 isolates across eight carbon sources, each graded 0-4; the mineral-medium control column was excluded from the niche profile. Presence/absence breadth is saturated in this collection — a mean of 7.33 of the eight sources, with 54 of 60 isolates using at least seven — so binarizing the scores would discard almost all between-strain variation, and the graded values were used throughout. Assimilation capacity was defined as the row sum of the graded scores (mean 12.1, range 2-23 of a possible 32).

Two questions were asked of these profiles. At the strain level, Spearman correlations related assimilation capacity to 18 network descriptors: the twelve layer-specific out-degrees, the four functional-category means, total out-degree and the participation coefficient, with Benjamini-Hochberg correction across the 18 tests. At the dyad level, a Mantel test related metabolic niche distance, taken as the Euclidean distance between graded assimilation profiles, to interaction presence within each layer, with Benjamini-Hochberg correction across the twelve layers. Neither analysis returned a significant association (Figure S11); the smallest corrected probability was $q = 0.085$, for the dyad-level test in the IC and IG layers. These analyses bound what carbohydrate assimilation predicts about network position and do not speak to metabolic properties the API panel does not capture.

**Supplementary Methods S10. Spatial configuration and paired analyses**

To assess spatial dependence, inhibition of growth (IG) and change in sporulation (CS) were evaluated under two configurations on the same plate as the direct-contact assay (Figure 1): a close-proximity condition in which the inocula were placed close enough for the colonies to expand into contact, and a spatially separated condition in which the inoculation points were approximately 10 mm apart (centre to centre). This design assessed the effect of spatial separation rather than distinguishing strictly between contact-dependent and diffusible mechanisms; reduced interaction frequencies under separation may involve contact-associated processes, diffusion or dilution of extracellular compounds, nutrient depletion, pH gradients or growth-dependent effects. The two configurations were scored as a separate paired dataset of four $60 \times 60$ binary matrices (IG and CS, close and separated), read from the same plates and readings as the direct-contact assay (Supplementary Methods S2) but with a spatial criterion, namely whether the response was present with the inocula in contact and, separately, with the inocula separated. The matrices share the strain order and the row/column convention of the twelve direct-contact layers (rows = the strain in which the phenotype is scored, columns = the partner). Because of the different criterion, the close-proximity IG and CS matrices differ from the IG and CS layers of Table 1 in 650 and 291 of 3,540 directed entries respectively; the spatial matrices are used only for the paired comparison and are never merged with the twelve-layer network.

Directed network density was calculated for each phenotype and configuration as the number of positive ordered non-self pairs divided by 3,540, and the spatial effect was expressed as $\Delta\rho = \rho_{close} - \rho_{separated}$. Each ordered pair was classified as persistent, lost after separation, gained after separation or absent under both configurations. Because the unordered strain dyad is the confrontation unit, all resampling moved both directed entries of a dyad together: the significance of $\Delta\rho$ was assessed against 10,000 paired randomizations that exchange the configuration label of a whole dyad, with $P = (b + 1)/(m + 1)$, and its 95% interval was obtained from 10,000 bootstrap resamples of dyads. McNemar's test and the exact binomial test on the discordant directed pairs (lost versus gained) are reported for comparison (IG: $\chi^2 = 501.1$, $P = 5.6 \times 10^{-111}$ and $P = 3.0 \times 10^{-141}$; CS: $\chi^2 = 0.76$, $P = 0.383$ and $P = 0.428$), and matched-pair odds ratios (lost:gained) are given with Wald 95% intervals on the log scale. These resamples are computational inference on the recorded matrices, not experimental replicates. Results are shown in Figure 6 and the matrices in Figure S12.

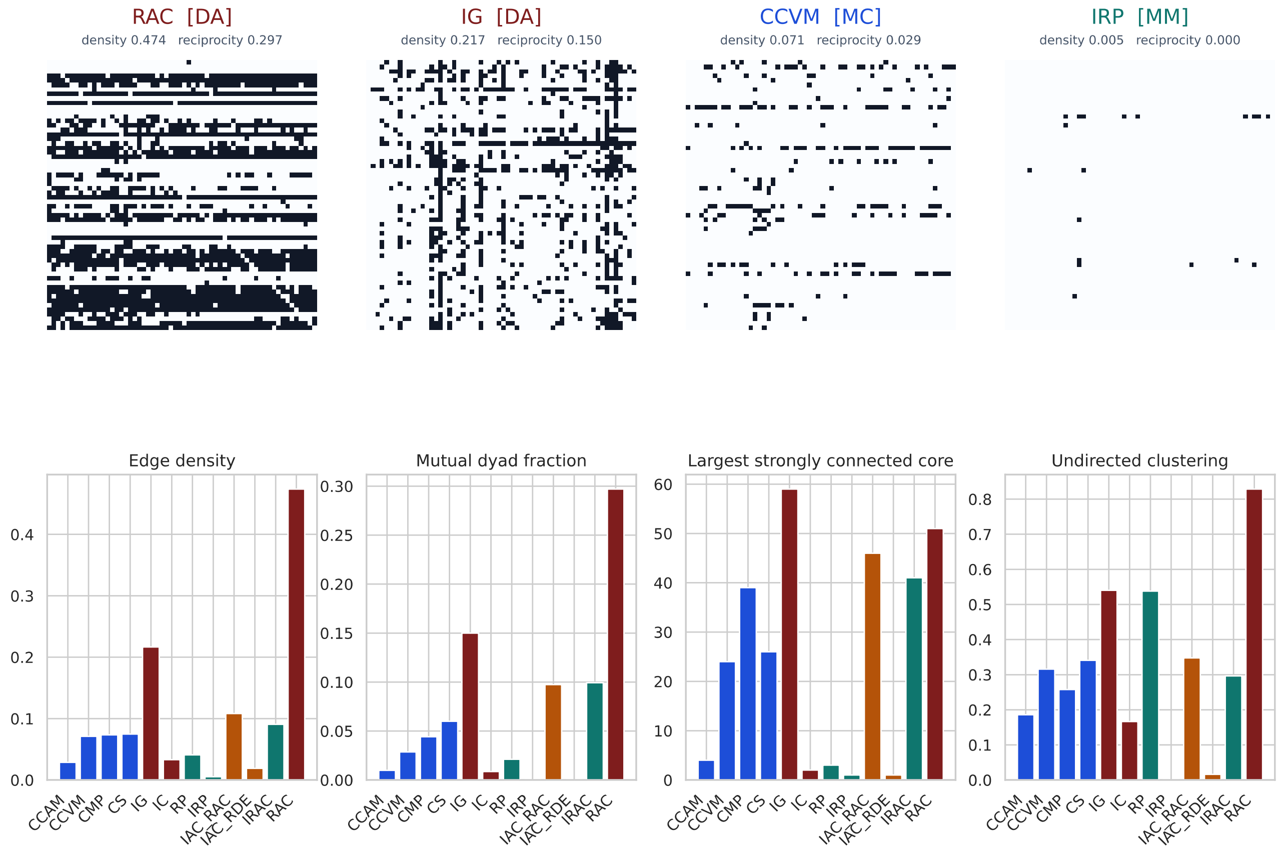


**Figure S1. Per-layer adjacency structure and layer-level summary.** Binary directed adjacency matrices for four layers (RAC, IG, CCVM and IRP) spanning the density range (rows=receivers, columns=senders); each panel gives that layer's directed edge density and pair-based reciprocity (Top row). All twelve layers compared on edge density, mutual dyad fraction, largest strongly connected component and undirected clustering, coloured by functional category. Direct-antagonism layers (RAC, IG) dominate density, reciprocity and SCC size, whereas developmental, morphogenetic and metabolic layers are sparser and more fragmented (Bottom row). Numerical values are given in Table 1.

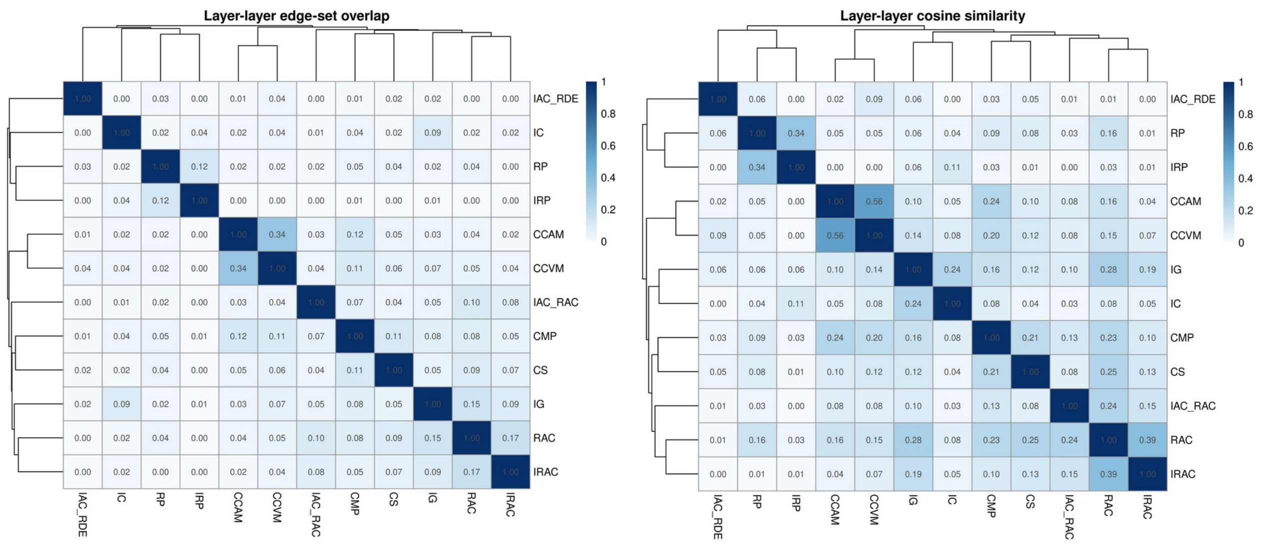


**Figure S2. Raw similarity among interaction layers, before comparison with null models.** Jaccard similarity of directed edge sets for all layer pairs (Left). Cosine similarity of the same layers. Both matrices are hierarchically clustered. Mean Jaccard similarity was 0.047 across the 66 layer pairs (Right). Because the expected Jaccard similarity depends on layer density, these raw values are descriptive only and are not evidence of layer independence; the degree-preserving comparison is shown in Figure 2.

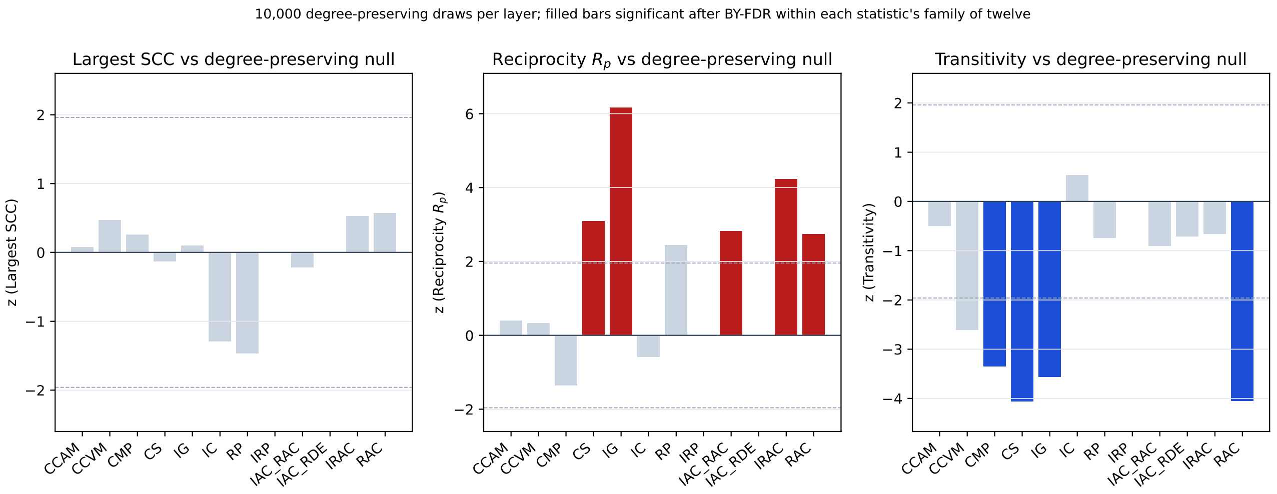


**Figure S3. Observed network structure relative to degree-preserving null models.** Standardized effect sizes (z) for the size of the largest strongly connected component (left), pair-based reciprocity (centre) and transitivity (right) for each interaction layer, against nulls generated by 10,000 degree-preserving directed edge-swap rewirings. Dashed lines mark $z$=±2. Largest-SCC sizes fall within null expectation in every layer, so the broad connectedness of IG and RAC follows from their degree distributions. Five layers are significantly more reciprocal than their degree sequences predict: IG most strongly ($z$=+6.17), then IRAC ($z$=+4.24), CS ($z$=+3.10), IAC_RAC ($z$=+2.82) and RAC ($z$=+2.75); none is significantly less reciprocal, and reciprocity is undefined for IRP and IAC_RDE, which contain no mutual dyad in any draw. Transitivity runs the other way: four layers are significantly less transitive than expected (CS $z$=-4.06, RAC $z$=-4.05, IG $z$=-3.56, CMP $z$=-3.34) and none more so, with CCVM close to the threshold (-2.60, $q$=0.064). Even dense, broadly connected layers are therefore not preferentially organised into closed triplets.

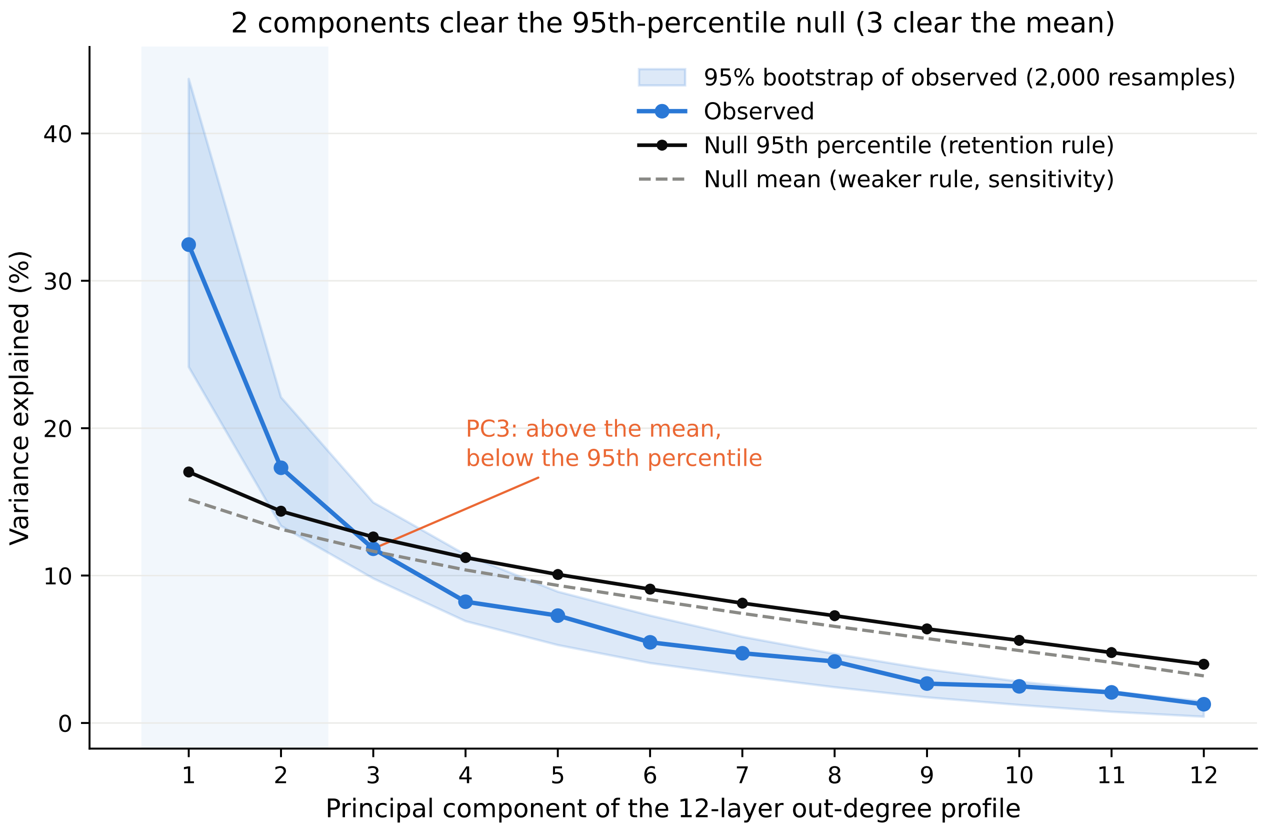


**Figure S4. Dimensionality of the twelve-layer out-degree profile by Horn's parallel analysis.** Observed variance explained by each principal component of the 12-dimensional outgoing profile, with a 95% interval from 2,000 nonparametric bootstrap resamples of the 60 strains, each re-standardized and re-decomposed. The null comes from 1,000 datasets in which values are permuted independently within each column, which preserves every layer's marginal distribution while destroying between-layer covariance. A component is retained when its observed variance exceeds the 95th percentile of that null (solid line); the mean-eigenvalue rule (dashed) is the weaker criterion reported alongside as a sensitivity analysis. PC1 (32.5%) and PC2 (17.3%) clear both. PC3 (11.8%) clears the mean (11.6%) but not the 95th percentile (12.6%), and its bootstrap interval spans the null, so it is reported as marginal rather than retained. PC4 (8.2%) falls below both.

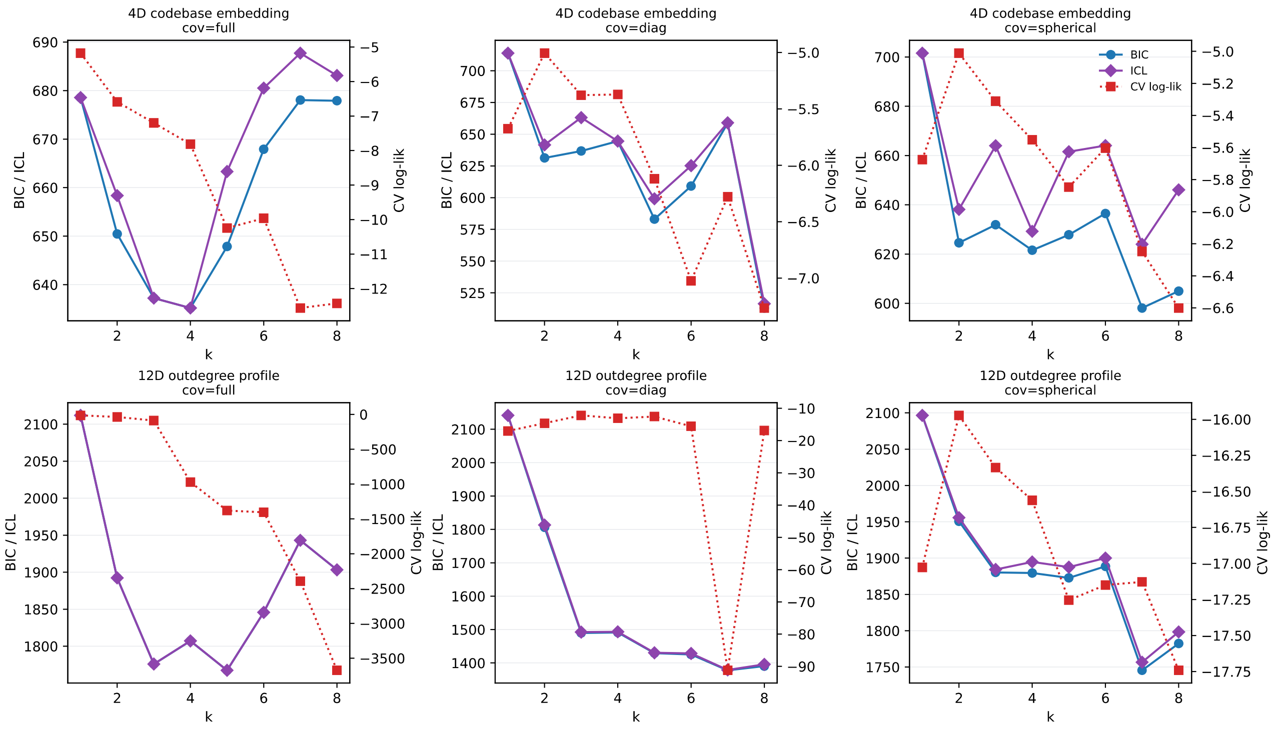


**Figure S5. Gaussian-mixture model-order selection gives no stable component count.** Mixture models were fitted for $k$=1-8 to the four-dimensional ecological embedding (top row) and the twelve-dimensional out-degree profile (bottom row) under full, diagonal and spherical covariance (columns). Every fit used 50 random initializations, a covariance regularization of $10^{-4}$, and a shared 10-fold partition, so all component counts are scored on identical splits. Left axis, BIC (circles) and ICL (diamonds), minimised at the preferred k; right axis, mean held-out log-likelihood. The information criteria prefer between four and eight components depending on the embedding and covariance structure, so they do not identify a single solution either. Free parameters grow steeply with $k$ under full covariance with 90 free parameters per component in the twelve-dimensional profile (78 covariance terms and 12 means), against 24 under diagonal and 13 under spherical, and the held-out likelihood of the full-covariance family degrades accordingly, from -14.53 at $k$=1 to -3673 at $k$=8. Those full-covariance solutions at $k$ > 3 are diagnostics of overfitting rather than candidate models. The highest held-out log-likelihood of any model examined was the diagonal three-component solution (-12.19), against -14.53 for the best full-covariance model and -15.97 for the best spherical one.

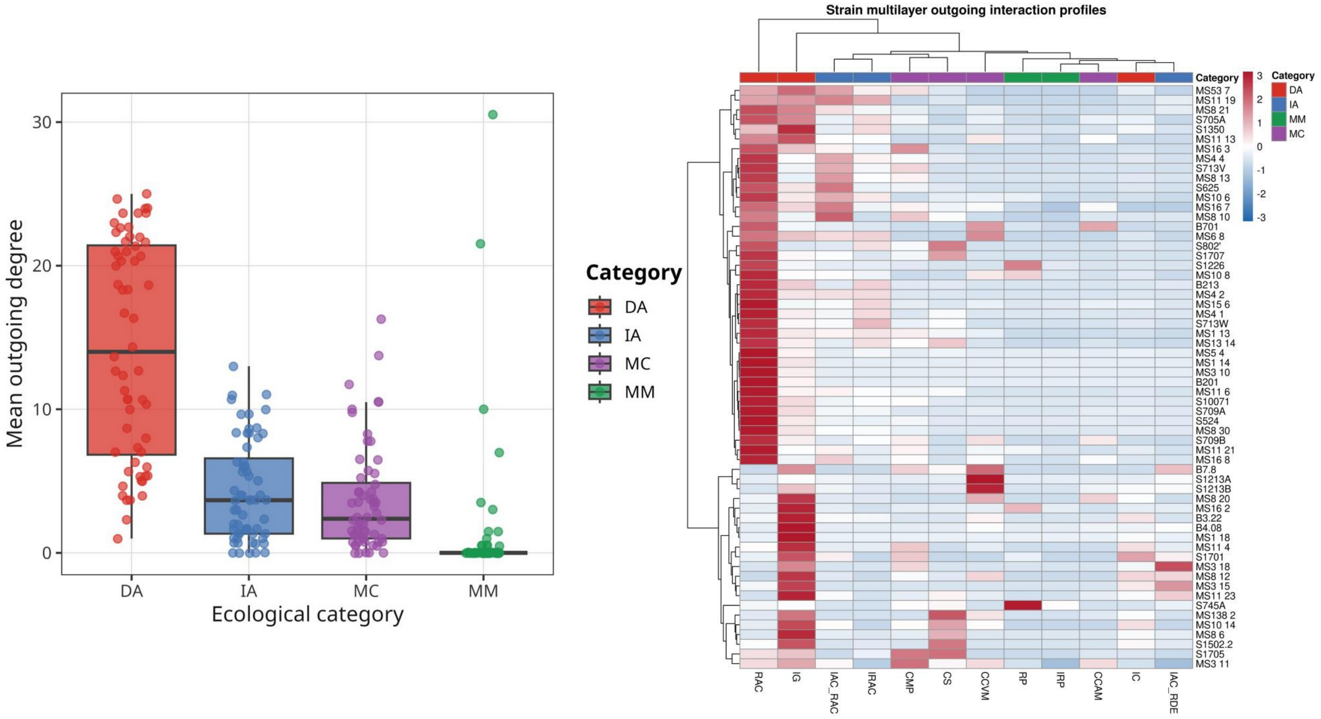


**Figure S6. Interaction output across functional categories and multilayer strain fingerprints.** Distribution of mean outgoing degree within each functional category (DA=direct antagonism, IA=induced antagonism, MC=morphogenetic control, MM=metabolic modulation); each point is one strain. Direct antagonism carries the highest average output, but within-category spread is wide, indicating multiple routes to community-level influence (Left). Hierarchically clustered heatmap of *z*-standardized outgoing interaction profiles across the twelve layers. Strains vary continuously across interaction dimensions and no sharply separated groups emerge (Right).

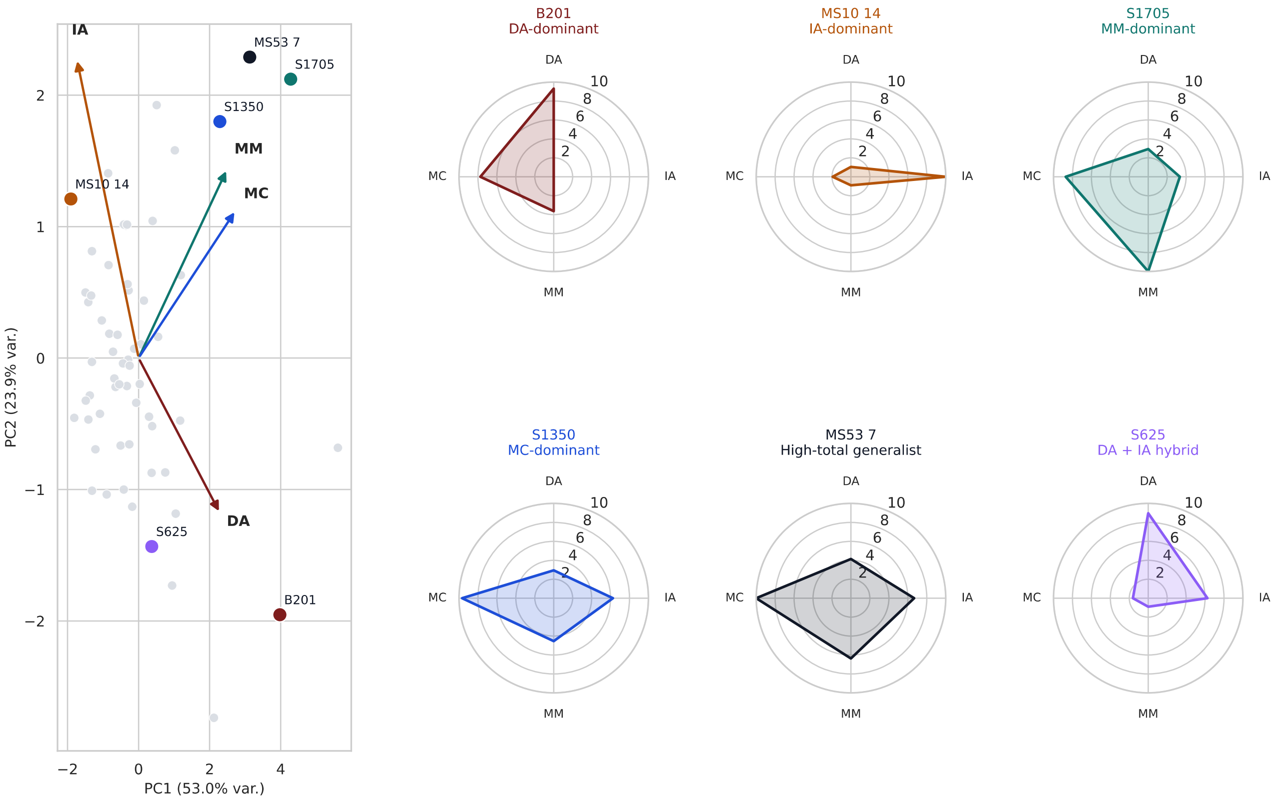


**Figure S7. Reduced four-dimensional embedding preserves the multilayer structure and illustrates the specialist-generalist continuum.** Principal component analysis of the four aggregated categories (DA, IA, MM, MC) for all strains; vectors give the ecological gradient defined by each category (Left). Radar plots for representative strains: B201 (DA-dominant), MS10-14 (IA-dominant), S1705 (MM-dominant), S1350 (MC-dominant), MS53-7 (broadly active) and S625 (DA-IA hybrid). The reduced embedding preserves the organisation of the twelve-layer space (Spearman $\rho$=0.934 between reduced and full PC1 scores) (Right). Because the four categories are defined a priori from the same twelve layers, this correspondence is a biologically interpretable compression rather than independent validation.

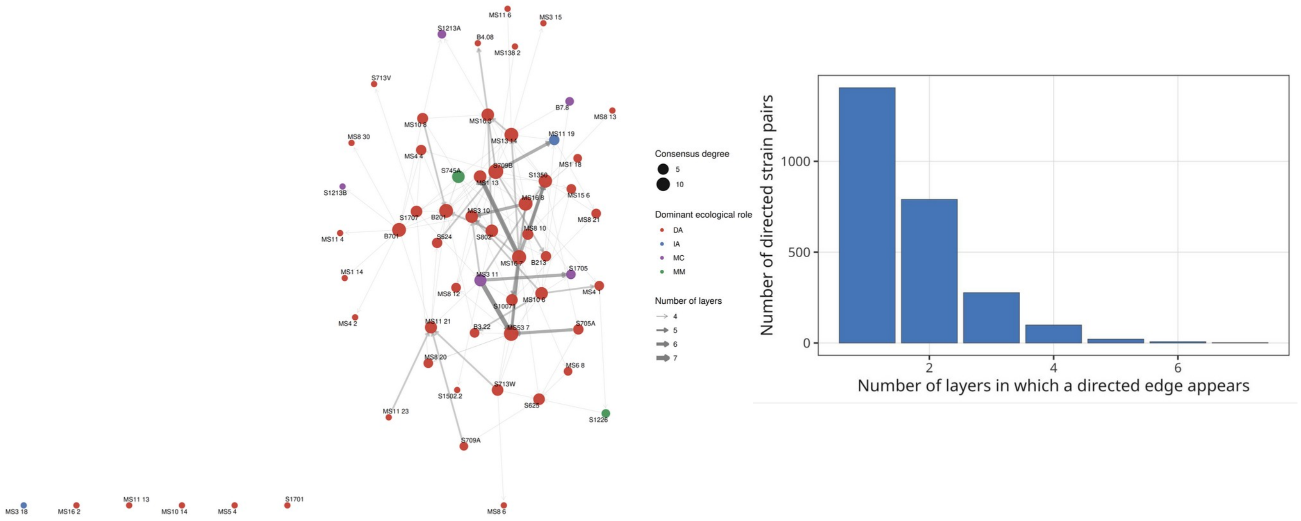


**Figure S8. Recurrent multilayer relationships are concentrated among highly interactive strains.** Consensus network retaining directed interactions present in at least four of the twelve layers; node size gives consensus degree, node colour the dominant ecological category and edge width the number of layers (Left). Distribution of the number of layers in which a directed strain pair appears. Most interactions occur in only one or two layers and recurrence across four or more is rare (Right). Because the ≥4-layer criterion is operational rather than biologically derived, the resulting topology should be read as heterogeneous multilayer recurrence rather than as demonstrated core-periphery organisation.

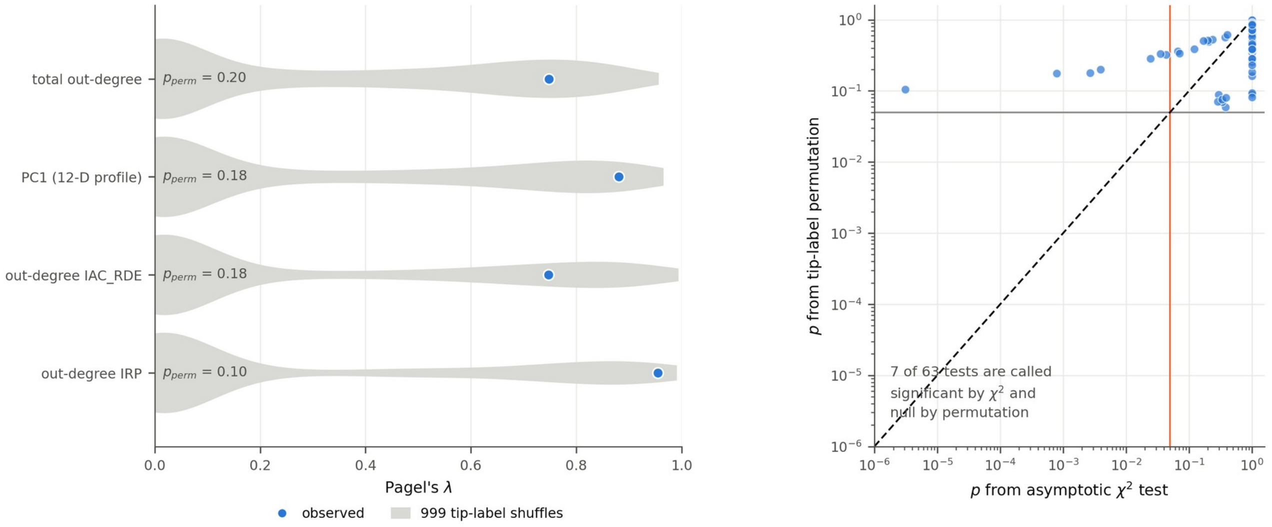


**Figure S9. Permutation testing removes spurious phylogenetic signal produced by asymptotic likelihood-ratio tests.** For the four traits with the smallest $\chi^2$ likelihood-ratio probabilities, the distribution of Pagel's λ under 999 tip-label shuffles with the observed λ marked; randomised trait values reproduce the observed estimate (Left). $\chi^2$ p-value against tip-label permutation p-value for all 63 trait x topology tests (log-log; dashed identity line, vertical line $\alpha$=0.05). All points lie above the identity and seven tests are called significant by $\chi^2$ but null by permutation. The cause is a rank-deficient phylogenetic covariance matrix (rank 44 of 60, from strains sharing identical 16S rRNA sequences) combined with a wide range in tip depth, which makes the asymptotic test anticonservative on these data. This panel is a calibration diagnostic for the testing procedure: it shows that the asymptotic likelihood-ratio test is anticonservative on this tree and that the permutation test is not. It does not by itself validate Pagel's λ as a model on a non-ultrametric gene tree, and the λ estimates are interpreted only against their own permutation nulls.

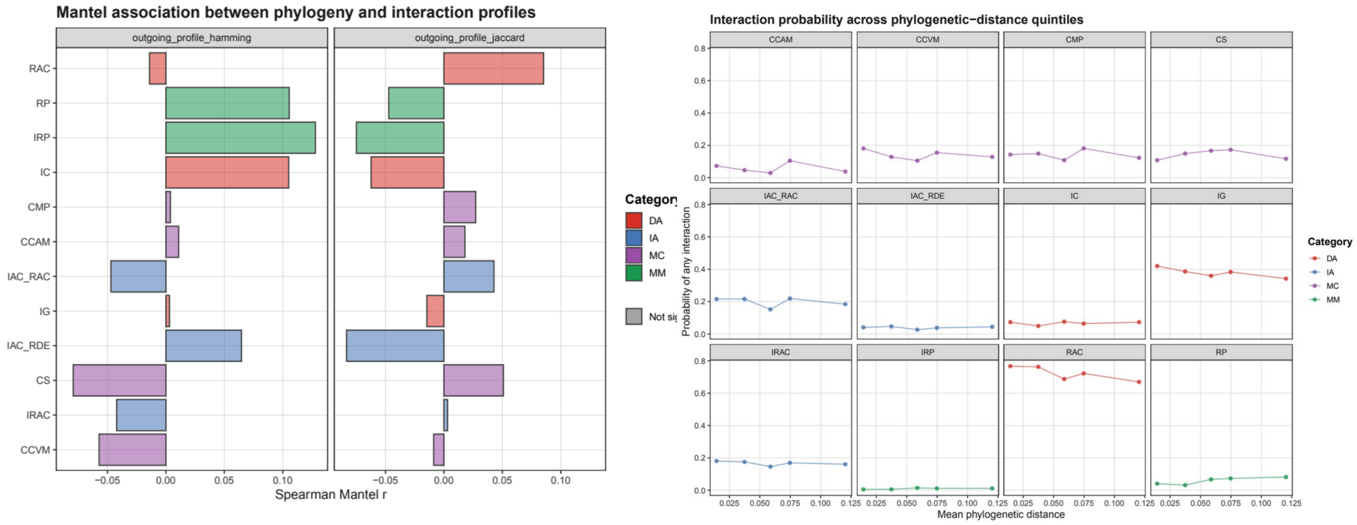


**Figure S10. Layer-specific relationships between 16S divergence and interaction profiles.** Spearman Mantel correlation between pairwise 16S rRNA genetic distance and layer-specific ecological distance, computed with Hamming (left) and Jaccard (right) profile distances; no layer reaches significance after correction. b, probability of a directed interaction in each layer across quintiles of increasing 16S rRNA phylogenetic distance. Interaction frequencies are broadly flat across evolutionary distance for antagonistic, induced-antagonistic, morphogenetic and metabolic phenotypes, so neither closely nor distantly related strains interact consistently more often.

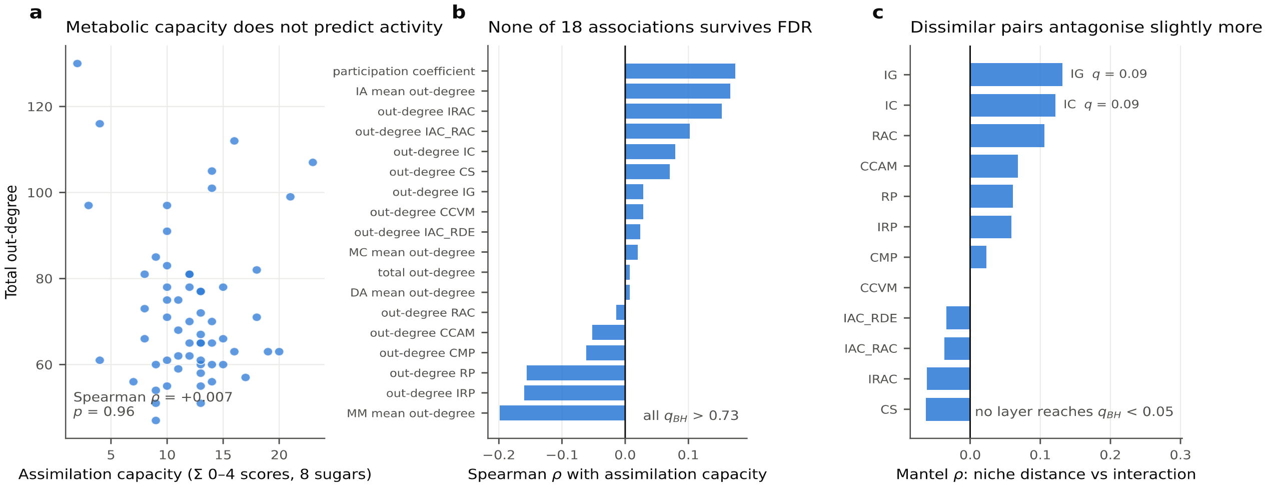


**Figure S11. Carbohydrate assimilation does not predict network position.** Graded carbohydrate assimilation capacity, the sum of 0-4 scores across eight carbon sources, against total out-degree (Spearman $\rho$=+0.007, $P$=0.96) (Left). All 18 strain-level associations between assimilation capacity and network descriptors, ranked; none reaches $q$ < 0.05 after Benjamini-Hochberg correction, and all exceed $q$=0.73 (Middle). Dyad-level Mantel correlation between metabolic niche distance and interaction presence within each layer; no layer is significant after correction, the smallest corrected probability being $q$=0.085 for IC and IG (Right). Methods are given in Supplementary Methods S9.

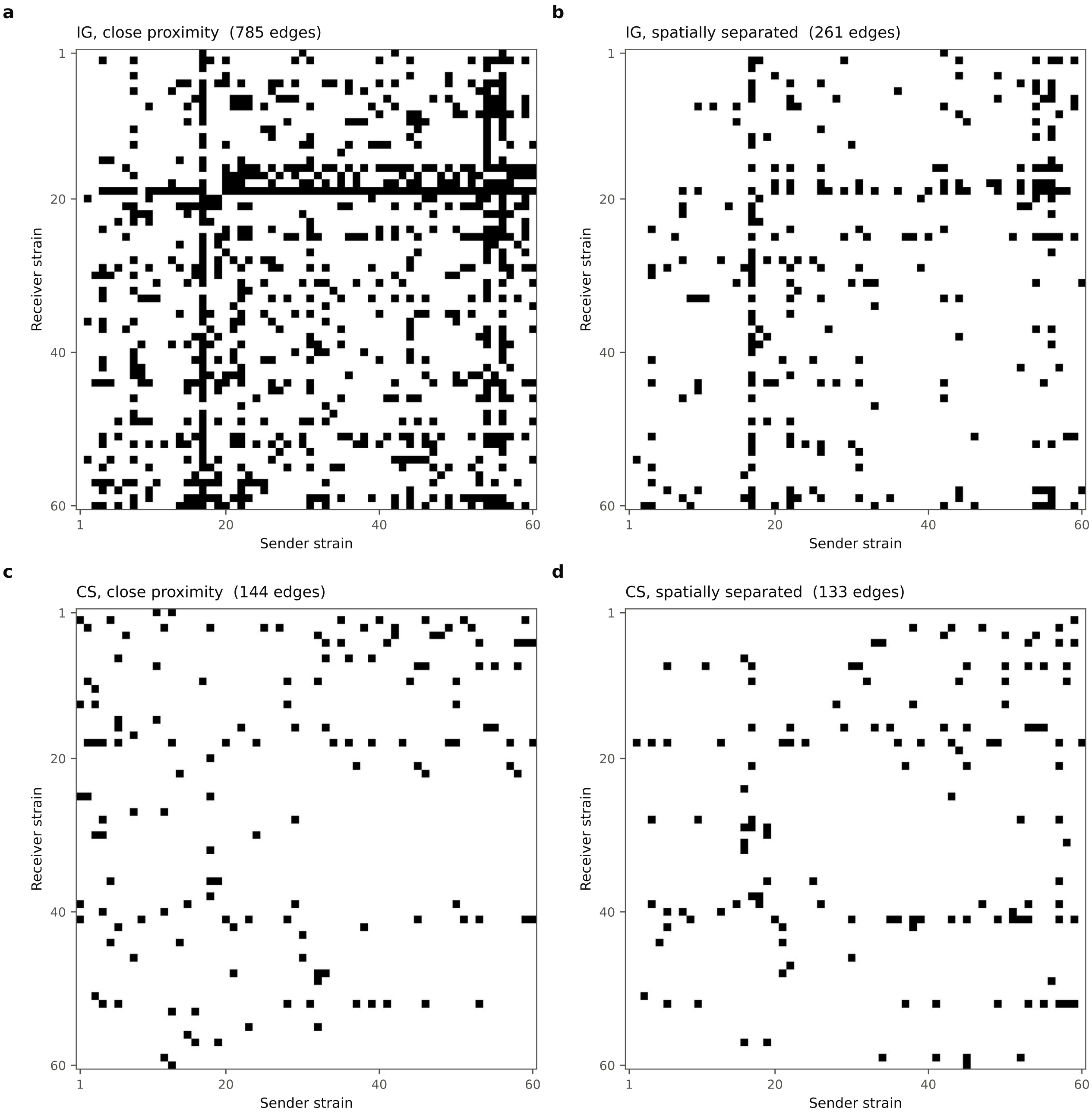


**Figure S12. Interaction matrices for growth inhibition and change in sporulation under both spatial configurations.** Binary directed matrices of the paired spatial experiment (rows=receivers, columns=senders, as in Figure S1; strains in the same order in all four panels). a, IG with the inocula in close proximity (785 edges); b, IG with the inocula spatially separated (261 edges); c, CS in close proximity (144 edges); d, CS spatially separated (133 edges). These matrices were scored separately from the twelve direct-contact layers and are used only for the paired comparison in Figure 6.

### Supplementary Tables

**Table S1.** Strain collection, 16S rRNA assignments and sequence accessions. Origin, taxonomic assignment, 16S rRNA gene accession numbers and sequence length for the 60 soil actinobacterial isolates, listed in order of sequence submission. Near-full-length 16S rRNA fragments were amplified with primers 27F/1492R, sequenced by Sanger chemistry and deposited in GenBank under the contiguous accession range PZ882263-PZ882322. Genus was assigned by 16S identity: 56 isolates are Streptomyces, one is Saccharothrix (MS53 7), two are Lentzea (MS10 8, MS3 15) and one is Amycolatopsis (MS3 18); the collection is described as actinobacterial throughout for that reason. One isolate appears as S1430 in the sequencing record and as S1226 in the interaction matrices; the two labels denote the same strain. Isolates are assigned to genus only; no closest-type-strain identity is reported. MS8 12 was isolated in September 2011 and the remaining isolates in September 2018. The raw sequencing file contains a second, shorter record labelled S705A (1,390 bp, 92% identical to the deposited sequence); it was not deposited and was not used, and PZ882302 is the S705A sequence used throughout.

| Strain | *Organism (16S rRNA assignment)* | GenBank accession | 16S rRNAlength (bp) | Isolation date | Geographical origin |
|---|---|---|---|---|---|
| S1226 | *Streptomyces sp.* | PZ882263 | 1485 | Sep-2018 | Oujda, Morocco |
| B201 | *Streptomyces sp.* | PZ882264 | 1376 | Sep-2018 | Fez, Teghat, Morocc[illegible] |
| MS53 7 | *Saccharothrix sp.* | PZ882265 | 1440 | Sep-2018 | Beni-Mellal, Morocc[illegible] |
| B213 | *Streptomyces sp.* | PZ882266 | 1418 | Sep-2018 | Fez, Teghat, Morocc[illegible] |
| B3.22 | *Streptomyces sp.* | PZ882267 | 1419 | Sep-2018 | Fez, Teghat, Morocc[illegible] |
| MS10 8 | *Lentzea sp.* | PZ882268 | 1409 | Sep-2018 | Agadir, Taghazout, M[illegible] |
| MS8 12 | *Streptomyces sp.* | PZ882269 | 1384 | Sep-2011 | Beni-Mellal, Morocc[illegible] |
| S1701 | *Streptomyces sp.* | PZ882270 | 1414 | Sep-2018 | Oujda, Morocco |
| MS11 4 | *Streptomyces sp.* | PZ882271 | 1447 | Sep-2018 | Agadir, Taghazout, M[illegible] |
| MS138 2 | *Streptomyces sp.* | PZ882272 | 1377 | Sep-2018 | Agadir, Taghazout, M[illegible] |
| S10071 | *Streptomyces sp.* | PZ882273 | 1457 | Sep-2018 | Oujda, Morocco |
| MS15 6 | *Streptomyces sp.* | PZ882274 | 1421 | Sep-2018 | Agadir, Taghazout, M[illegible] |
| MS6 8 | *Streptomyces sp.* | PZ882275 | 1370 | Sep-2018 | Beni-Mellal, Morocc[illegible] |
| MS8 20 | *Streptomyces sp.* | PZ882276 | 1478 | Sep-2018 | Beni-Mellal, Morocc[illegible] |
| MS3 11 | *Streptomyces sp.* | PZ882277 | 1379 | Sep-2018 | Agadir, Taghazout, M[illegible] |
| S625 | *Streptomyces sp.* | PZ882278 | 1389 | Sep-2018 | Saïdia, Morocco |
| MS3 10 | *Streptomyces sp.* | PZ882279 | 1448 | Sep-2018 | Agadir, Taghazout, M[illegible] |
| MS1 14 | *Streptomyces sp.* | PZ882280 | 1378 | Sep-2018 | Agadir, Taghazout, M[illegible] |
| MS11 21 | *Streptomyces sp.* | PZ882281 | 1395 | Sep-2018 | Agadir, Taghazout, M[illegible] |
| MS11 6 | *Streptomyces sp.* | PZ882282 | 1359 | Sep-2018 | Agadir, Taghazout, M[illegible] |
| MS8 13 | *Streptomyces sp.* | PZ882283 | 1441 | Sep-2018 | Beni-Mellal, Morocc[illegible] |
| MS1 13 | *Streptomyces sp.* | PZ882284 | 1424 | Sep-2018 | Agadir, Taghazout, M[illegible] |
| MS16 3 | *Streptomyces sp.* | PZ882285 | 1445 | Sep-2018 | Agadir, Taghazout, M[illegible] |
| S1705 | *Streptomyces sp.* | PZ882286 | 1388 | Sep-2018 | Oujda, Morocco |
| S524 | *Streptomyces sp.* | PZ882287 | 1389 | Sep-2018 | Saïdia, Morocco |
| B701 | *Streptomyces sp.* | PZ882288 | 1431 | Sep-2018 | Fez, Teghat, Morocc[illegible] |
| MS11 13 | *Streptomyces sp.* | PZ882289 | 1343 | Sep-2018 | Agadir, Taghazout, M[illegible] |
| MS4 1 | *Streptomyces sp.* | PZ882290 | 1447 | Sep-2018 | Beni-Mellal, Morocc[illegible] |
| S1213A | *Streptomyces sp.* | PZ882291 | 1444 | Sep-2018 | Oujda, Morocco |
| S709A | *Streptomyces sp.* | PZ882292 | 1385 | Sep-2018 | Saïdia, Morocco |
| B4.08 | *Streptomyces sp.* | PZ882293 | 1494 | Sep-2018 | Fez, Teghat, Morocc[illegible] |
| B7.8 | *Streptomyces sp.* | PZ882294 | 1501 | Sep-2018 | Fez, Teghat, Morocc[illegible] |
| MS10 14 | *Streptomyces sp.* | PZ882295 | 1409 | Sep-2018 | Agadir, Taghazout, M[illegible] |
| MS11 19 | *Streptomyces sp.* | PZ882296 | 1387 | Sep-2018 | Agadir, Taghazout, M[illegible] |
| MS13 14 | *Streptomyces sp.* | PZ882297 | 1316 | Sep-2018 | Agadir, Taghazout, M[illegible] |
| MS16 7 | *Streptomyces sp.* | PZ882298 | 1390 | Sep-2018 | Agadir, Taghazout, M[illegible] |
| MS16 8 | *Streptomyces sp.* | PZ882299 | 1469 | Sep-2018 | Agadir, Taghazout, M[illegible] |
| MS3 15 | *Lentzea sp.* | PZ882300 | 1393 | Sep-2018 | Beni-Mellal, Morocc[illegible] |
| MS3 18 | *Amycolatopsis sp.* | PZ882301 | 1426 | Sep-2018 | Beni-Mellal, Morocc[illegible] |

| | | | | | |
|---|---|---|---|---|---|
| S705A | *Streptomyces sp.* | PZ882302 | 1468 | Sep-2018 | Saïdia, Morocco |
| MS1 18 | *Streptomyces sp.* | PZ882303 | 1492 | Sep-2018 | Agadir, Taghazout, M |
| MS8 6 | *Streptomyces sp.* | PZ882304 | 1430 | Sep-2018 | Beni-Mellal, Morocc |
| MS8 21 | *Streptomyces sp.* | PZ882305 | 1417 | Sep-2018 | Beni-Mellal, Morocc |
| MS8 30 | *Streptomyces sp.* | PZ882306 | 1460 | Sep-2018 | Beni-Mellal, Morocc |
| S713'V | *Streptomyces sp.* | PZ882307 | 1436 | Sep-2018 | Saïdia, Morocco |
| S1213B | *Streptomyces sp.* | PZ882308 | 1461 | Sep-2018 | Oujda, Morocco |
| S802' | *Streptomyces sp.* | PZ882309 | 1476 | Sep-2018 | Saïdia, Morocco |
| MS4 4 | *Streptomyces sp.* | PZ882310 | 1447 | Sep-2018 | Beni-Mellal, Morocc |
| S1707 | *Streptomyces sp.* | PZ882311 | 1422 | Sep-2018 | Oujda, Morocco |
| S713'W | *Streptomyces sp.* | PZ882312 | 1463 | Sep-2018 | Saïdia, Morocco |
| MS4 2 | *Streptomyces sp.* | PZ882313 | 1478 | Sep-2018 | Beni-Mellal, Morocc |
| S745A | *Streptomyces sp.* | PZ882314 | 1415 | Sep-2018 | Saïdia, Morocco |
| S1502.2 | *Streptomyces sp.* | PZ882315 | 1476 | Sep-2018 | Oujda, Morocco |
| MS5 4 | *Streptomyces sp.* | PZ882316 | 1442 | Sep-2018 | Beni-Mellal, Morocc |
| MS11 23 | *Streptomyces sp.* | PZ882317 | 1462 | Sep-2018 | Agadir, Taghazout, M |
| MS10 6 | *Streptomyces sp.* | PZ882318 | 1426 | Sep-2018 | Agadir, Taghazout, M |
| S709B | *Streptomyces sp.* | PZ882319 | 1420 | Sep-2018 | Saïdia, Morocco |
| S1350 | *Streptomyces sp.* | PZ882320 | 1439 | Sep-2018 | Oujda, Morocco |
| MS8 10 | *Streptomyces sp.* | PZ882321 | 1376 | Sep-2018 | Beni-Mellal, Morocc |
| MS16 2 | *Streptomyces sp.* | PZ882322 | 1417 | Sep-2018 | Agadir, Taghazout, M |

**Table S2.**

Cross-layer overlap for all 66 layer pairs relative to degree-preserving nulls. All 66 layer pairs. Each layer was randomized with 10,000 degree-preserving directed edge-swap permutations retaining its empirical in- and out-degree sequences (seed 20260522). Two-sided empirical probabilities use the Phipson-Smyth $(M+1)/(N+1)$ correction and are corrected across all 66 pairs by both Benjamini-Hochberg and Benjamini-Yekutieli; BY is used for inference because each layer appears in eleven of the tests. This table and Table 2 are generated from the same results object, so their *z* values agree by construction.

| Layer 1 | Layer 2 | *J* observed | *J* null mean | *J* null SD | *z* | *p* (empirical) | *q* (BH) | *q* (BY) | Significant (BY) | Shared assay co |
|---|---|---|---|---|---|---|---|---|---|---|
| CCAM | CCVM | 0.3358 | 0.1292 | 0.0128 | 16.19 | 0.0001 | 0.0009 | 0.0045 | yes | yes |
| CCAM | CMP | 0.1207 | 0.053 | 0.0088 | 7.69 | 0.0001 | 0.0009 | 0.0045 | yes | no |
| CCAM | CS | 0.0456 | 0.0226 | 0.006 | 3.86 | 0.0006 | 0.004 | 0.0189 | yes | no |
| CCAM | IG | 0.0345 | 0.0307 | 0.0038 | 0.99 | 0.3425 | 0.6625 | 1 | no | no |
| CCAM | IC | 0.0234 | 0.0144 | 0.0061 | 1.45 | 0.2373 | 0.58 | 1 | no | no |
| CCAM | RP | 0.0249 | 0.0251 | 0.006 | -0.04 | 1 | 1 | 1 | no | no |
| CCAM | IRP | 0 | 0.0017 | 0.0036 | -0.47 | 1 | 1 | 1 | no | no |
| CCAM | IAC_RAC | 0.0341 | 0.0285 | 0.006 | 0.94 | 0.3655 | 0.6625 | 1 | no | no |
| CCAM | IAC_RDE | 0.012 | 0.009 | 0.0049 | 0.6 | 1 | 1 | 1 | no | no |
| CCAM | IRAC | 0.0193 | 0.0219 | 0.0059 | -0.45 | 0.6841 | 0.8886 | 1 | no | no |
| CCAM | RAC | 0.0385 | 0.0378 | 0.0018 | 0.38 | 0.7389 | 0.9202 | 1 | no | no |
| CCVM | CMP | 0.1082 | 0.0473 | 0.0074 | 8.23 | 0.0001 | 0.0009 | 0.0045 | yes | no |
| CCVM | CS | 0.0616 | 0.0254 | 0.0057 | 6.41 | 0.0001 | 0.0009 | 0.0045 | yes | no |
| CCVM | IG | 0.0659 | 0.0611 | 0.0051 | 0.94 | 0.3815 | 0.6625 | 1 | no | no |
| CCVM | IC | 0.0394 | 0.0274 | 0.0063 | 1.91 | 0.0698 | 0.3071 | 1 | no | no |
| CCVM | RP | 0.0232 | 0.0247 | 0.0056 | -0.27 | 0.8192 | 1 | 1 | no | no |
| CCVM | IRP | 0 | 0.0046 | 0.0035 | -1.29 | 0.3249 | 0.6625 | 1 | no | no |
| CCVM | IAC_RAC | 0.0393 | 0.0428 | 0.0065 | -0.54 | 0.6034 | 0.8297 | 1 | no | no |
| CCVM | IAC_RDE | 0.0391 | 0.0294 | 0.0053 | 1.82 | 0.1114 | 0.3768 | 1 | no | no |
| CCVM | IRAC | 0.038 | 0.0371 | 0.0065 | 0.14 | 1 | 1 | 1 | no | no |
| CCVM | RAC | 0.0524 | 0.0543 | 0.0023 | -0.83 | 0.4536 | 0.7484 | 1 | no | no |
| CMP | CS | 0.1146 | 0.081 | 0.0091 | 3.68 | 0.0006 | 0.004 | 0.0189 | yes | no |
| CMP | IG | 0.0754 | 0.0701 | 0.0057 | 0.93 | 0.3771 | 0.6625 | 1 | no | no |
| CMP | IC | 0.0386 | 0.0346 | 0.0074 | 0.53 | 0.6941 | 0.8886 | 1 | no | no |
| CMP | RP | 0.0465 | 0.0374 | 0.0051 | 1.78 | 0.1142 | 0.3768 | 1 | no | no |
| CMP | IRP | 0.0072 | 0.0082 | 0.0044 | -0.22 | 1 | 1 | 1 | no | no |
| CMP | IAC_RAC | 0.0681 | 0.0604 | 0.008 | 0.97 | 0.3782 | 0.6625 | 1 | no | no |
| CMP | IAC_RDE | 0.0124 | 0.0163 | 0.0055 | -0.71 | 0.5734 | 0.8171 | 1 | no | no |
| CMP | IRAC | 0.0544 | 0.0487 | 0.0077 | 0.75 | 0.4672 | 0.752 | 1 | no | no |
| CMP | RAC | 0.0833 | 0.0813 | 0.0029 | 0.69 | 0.5317 | 0.7799 | 1 | no | no |
| CS | IG | 0.0541 | 0.0609 | 0.0051 | -1.32 | 0.1986 | 0.5698 | 1 | no | no |
| CS | IC | 0.0187 | 0.0281 | 0.0062 | -1.54 | 0.1717 | 0.5396 | 1 | no | no |
| CS | RP | 0.0406 | 0.0297 | 0.0043 | 2.52 | 0.0169 | 0.0929 | 0.4437 | no | no |
| CS | IRP | 0.0035 | 0.0077 | 0.0041 | -1.03 | 0.4044 | 0.6843 | 1 | no | no |
| CS | IAC_RAC | 0.0385 | 0.0429 | 0.0063 | -0.71 | 0.5041 | 0.766 | 1 | no | no |
| CS | IAC_RDE | 0.0184 | 0.0129 | 0.0048 | 1.15 | 0.3266 | 0.6625 | 1 | no | no |
| CS | IRAC | 0.0674 | 0.0704 | 0.0076 | -0.4 | 0.7001 | 0.8886 | 1 | no | no |
| CS | RAC | 0.0935 | 0.0855 | 0.0022 | 3.67 | 0.0004 | 0.0033 | 0.0158 | yes | no |
| IG | IC | 0.09 | 0.0775 | 0.0048 | 2.59 | 0.0108 | 0.0648 | 0.3094 | no | no |
| IG | RP | 0.0213 | 0.0284 | 0.0031 | -2.29 | 0.0237 | 0.1203 | 0.5744 | no | no |
| IG | IRP | 0.009 | 0.0095 | 0.0019 | -0.24 | 1 | 1 | 1 | no | no |
| IG | IAC_RAC | 0.0474 | 0.0569 | 0.0053 | -1.81 | 0.0807 | 0.3164 | 1 | no | no |
| IG | IAC_RDE | 0.0171 | 0.0206 | 0.0035 | -1.02 | 0.3692 | 0.6625 | 1 | no | no |
| IG | IRAC | 0,0924 | 0,0648 | 0,0057 | 4,82 | 0,0001 | 0,0009 | 0,0045 | yes | no |
| IG | RAC | 0,1469 | 0,1431 | 0,0033 | 1,14 | 0,2872 | 0,6536 | 1 | no | no |
| IC | RP | 0,0195 | 0,017 | 0,0045 | 0,54 | 0,662 | 0,8886 | 1 | no | no |
| IC | IRP | 0,0382 | 0,0321 | 0,0034 | 1,81 | 0,2215 | 0,58 | 1 | no | no |

| | | | | | | | | | | |
|---|---|---|---|---|---|---|---|---|---|---|
| IC | IAC_RAC | 0,0121 | 0,0175 | 0,0047 | -1,13 | 0,2867 | 0,6536 | 1 | no | no |
| IC | IAC_RDE | 0 | 0,0006 | 0,0018 | -0,35 | 1 | 1 | 1 | no | no |
| IC | IRAC | 0,021 | 0,0206 | 0,0056 | 0,06 | 1 | 1 | 1 | no | no |
| IC | RAC | 0,0199 | 0,022 | 0,0017 | -1,25 | 0,2329 | 0,58 | 1 | no | no |
| RP | IRP | 0,1233 | 0,1052 | 0,0095 | 1,91 | 0,1135 | 0,3768 | 1 | no | no |
| RP | IAC_RAC | 0,0154 | 0,0188 | 0,0045 | -0,76 | 0,5106 | 0,766 | 1 | no | no |
| RP | IAC_RDE | 0,0291 | 0,0278 | 0,0025 | 0,51 | 1 | 1 | 1 | no | no |
| RP | IRAC | 0,0043 | 0,0119 | 0,0039 | -1,97 | 0,0815 | 0,3164 | 1 | no | no |
| RP | RAC | 0,0441 | 0,0421 | 0,0015 | 1,33 | 0,231 | 0,58 | 1 | no | no |
| IRP | IAC_RAC | 0 | 0,002 | 0,002 | -1,02 | 0,5818 | 0,8171 | 1 | no | no |
| IRP | IAC_RDE | 0 | 0 | 0 | +nan | 1 | 1 | 1 | no | no |
| IRP | IRAC | 0,0029 | 0,0024 | 0,0022 | 0,27 | 1 | 1 | 1 | no | no |
| IRP | RAC | 0,0036 | 0,0042 | 0,0008 | -0,77 | 0,4793 | 0,7531 | 1 | no | no |
| IAC_RAC | IAC_RDE | 0,0045 | 0,0054 | 0,003 | -0,3 | 1 | 1 | 1 | no | no |
| IAC_RAC | IRAC | 0,0781 | 0,087 | 0,0093 | -0,95 | 0,3625 | 0,6625 | 1 | no | no |
| IAC_RAC | RAC | 0,104 | 0,1247 | 0,0039 | -5,32 | 0,0001 | 0,0009 | 0,0045 | yes | no |
| IAC_RDE | IRAC | 0 | 0,005 | 0,0031 | -1,63 | 0,1842 | 0,5525 | 1 | no | no |
| IAC_RDE | RAC | 0,0023 | 0,0041 | 0,0009 | -1,97 | 0,0629 | 0,2965 | 1 | no | no |
| IRAC | RAC | 0,1664 | 0,1331 | 0,0036 | 9,24 | 0,0001 | 0,0009 | 0,0045 | yes | no |

**Table S3.** Concordance between two estimators of layer-specific ecological dominance. Normalized David's scores and Bradley-Terry strengths were estimated independently for every isolate within each of the twelve interaction layers. Within-layer Spearman correlations range from 0.59 to 0.99 (mean 0.87), so the weak agreement of strain rankings between layers (mean $\rho$=0.063) reflects phenotype-specific ecological roles rather than instability of the estimator. Because both measures derive from the same interaction matrices, their agreement is methodological concordance and not independent validation.

| **Layer** | **Spearman $\rho$ (David's score vs Bradley-Terry strength)** | ***P*** |
|---|---|---|
| CCAM | 0.834 | $1.32*10^{-16}$ |
| CCVM | 0.845 | $2.09*10^{-17}$ |
| CMP | 0.844 | $2.46*10^{-17}$ |
| CS | 0.903 | $6.29*10^{-23}$ |
| IG | 0.964 | $4.39*10^{-35}$ |
| IC | 0.588 | $7.74*10^{-7}$ |
| RP | 0.834 | $1.27*10^{-16}$ |
| IRP | 0.994 | $6.80*10^{-58}$ |
| IAC_RAC | 0.863 | $7.41*10^{-19}$ |
| IAC_RDE | 0.895 | $5.63*10^{-22}$ |
| IRAC | 0.892 | $1.24*10^{-22}$ |
| RAC | 0.973 | $1.09*10^{-38}$ |

**Table S4.** Taxonomic sensitivity analysis. Headline analyses repeated after excluding the four non-Streptomyces isolates (MS53 7, MS10 8, MS3 15, MS3 18) and compared with the complete 60-isolate cohort. Both arms now use 10,000 null draws, so the comparison carries no draw-count effect; at the matched count the full cohort returns the same eight over-overlapping pairs as the published analysis. Two pairs, CCAM-CS and CMP-CS, fall below the FDR threshold at *N*=56 and no new pair rises above it, which is the whole of the change. Layer-density and largest-SCC rankings are essentially unchanged, two principal components remain above the 95th-percentile parallel-analysis null in both cohorts, and cross-validated mixture modelling continues to favour a single component under full covariance. No conclusion in the manuscript depends on the four excluded isolates.

| **Quantity** | **Full cohort (*N* = 60)** | **Streptomyces only (*N* = 56)** |
|---|---|---|
| Jaccard null: over-overlapping pairs, published 10,000 draws | 8 | - |
| Jaccard null: over-overlapping pairs, matched 10,000 draws | 8 | 6 |
| Jaccard null: under-overlapping pairs, matched 10,000 draws | 1 | 1 |
| 12-D PCA PC1 (% variance) | 32.5 | 31.2 |
| 12-D PCA PC2 (% variance) | 17.3 | 15.2 |
| PCs above Horn parallel-analysis null | 2 | 2 |
| GMM components by 10-fold CV log-likelihood (12-D, full cov) | 1 | 1 |
| Layer density rank correlation *N*=60 vs *N*=56 (Spearman) | - | 1 |
| Largest-SCC rank correlation n60 vs n56 (Spearman) | - | 0.9982 |

### Supplementary references


1. Katoh K, Standley DM. MAFFT multiple sequence alignment software version 7: improvements in performance and usability. Mol Biol Evol 2013;30:772-80. https://doi.org/10.1093/molbev/mst010
2. Capella-Gutiérrez S, Silla-Martínez JM, Gabaldón T. trimAl: a tool for automated alignment trimming in large-scale phylogenetic analyses. Bioinformatics 2009;25:1972-3. https://doi.org/10.1093/bioinformatics/btp348
3. Price MN, Dehal PS, Arkin AP. FastTree 2-approximately maximum-likelihood trees for large alignments. PLoS ONE 2010;5:e9490. https://doi.org/10.1371/journal.pone.0009490
4. Piñeiro C, Abuín JM, Pichel JC. Very Fast Tree: speeding up the estimation of phylogenies for large alignments through parallelization and vectorization strategies. Bioinformatics 2020;36:4658-9. https://doi.org/10.1093/bioinformatics/btaa582
5. Davelos AL, Kinkel LL, Samac DA. Spatial variation in frequency and intensity of antibiotic interactions among streptomycetes from prairie soil. Appl Environ Microbiol 2004;70:1051-8. https://doi.org/10.1128/AEM.70.2.1051-1058.2004
6. Tarjan R. Depth-first search and linear graph algorithms. SIAM J Comput 1972;1:146-60. https://doi.org/10.1137/0201010
7. Guimerà R, Amaral LAN. Functional cartography of complex metabolic networks. Nature 2005;433:895-900. https://doi.org/10.1038/nature03288
8. Jaccard P. The distribution of the flora in the alpine zone. New Phytol 1912;11:37-50. https://doi.org/10.1111/j.1469-8137.1912.tb05611.x
9. Maslov S, Sneppen K. Specificity and stability in topology of protein networks. Science 2002;296:910-13. https://doi.org/10.1126/science.1065103
10. Phipson B, Smyth GK. Permutation P-values should never be zero: calculating exact P-values when permutations are randomly drawn. Stat Appl Genet Mol Biol 2010;9:Article39. https://doi.org/10.2202/1544-6115.1585
11. Benjamini Y, Yekutieli D. The control of the false discovery rate in multiple testing under dependency. Ann Stat 2001;29:1165-88. https://doi.org/10.1214/aos/1013699998
12. Benjamini Y, Hochberg Y. Controlling the false discovery rate: a practical and powerful approach to multiple testing. J R Stat Soc Series B Stat Methodol 1995;57:289-300. https://doi.org/10.1111/j.2517-6161.1995.tb02031.x
13. Pedregosa F, Varoquaux G, Gramfort A et al. Scikit-learn: machine learning in Python. J Mach Learn Res 2011;12:2825-30.
14. Horn JL. A rationale and test for the number of factors in factor analysis. Psychometrika 1965;30:179-85. https://doi.org/10.1007/BF02289447

15. Celeux G, Govaert G. Gaussian parsimonious clustering models. Pattern Recognit 1995;28:781-93. https://doi.org/10.1016/0031-3203(94)00125-6
16. Smyth P. Model selection for probabilistic clustering using cross-validated likelihood. Stat Comput 2000;10:63-72. https://doi.org/10.1023/A:1008940618127
17. Hartigan JA, Hartigan PM. The dip test of unimodality. Ann Stat 1985;13:70-84. https://doi.org/10.1214/aos/1176346577
18. David HA. Ranking from unbalanced paired-comparison data. Biometrika 1987;74:432-6. https://doi.org/10.1093/biomet/74.2.432
19. de Vries H, Stevens JMG, Vervaecke H. Measuring and testing the steepness of dominance hierarchies. Anim Behav 2006;71:585-92. https://doi.org/10.1016/j.anbehav.2005.05.015
20. Bradley RA, Terry ME. Rank analysis of incomplete block designs: I. The method of paired comparisons. Biometrika 1952;39:324-45. https://doi.org/10.1093/biomet/39.3-4.324
21. Pagel M. Inferring the historical patterns of biological evolution. Nature 1999;401:877-84. https://doi.org/10.1038/44766
22. Mantel N. The detection of disease clustering and a generalized regression approach. Cancer Res 1967;27:209-20.
